# Fibroblastic FLT3L supports lymph node dendritic cells in the interfollicular niche

**DOI:** 10.1101/2024.08.12.607692

**Authors:** Ryan S. Lane, Sunny Z. Wu, Christopher Davidson, Ashley Byrne, Brandon D. Kayser, Hejin Huang, Katherine Williams, Matthew Fernandez, Jian Jiang, Juan Zhang, Raymond Asuncion, Jérémie Decalf, Merone Roose-Girma, Wyne P. Lee, Lisa McGinnis, Soren Warming, William Stephenson, Sandra Rost, Christine Moussion, Tommaso Biancalani, Sören Müller, Shannon J. Turley

## Abstract

Dendritic cell (DC) homeostasis is maintained in secondary lymphoid organs (SLOs) by Fms-like tyrosine kinase 3 ligand (FLT3L). The specific niche providing this DC growth factor within human and mouse SLOs is unclear. Here, we show that Gremlin1 (Grem1)-expressing lymph node fibroblastic reticular cells (FRCs) support DC homeostasis via provision of FLT3L. Grem1 FRCs co-localize with DCs and express FLT3L in human and mouse lymph nodes. Using a new genetic model, we provide evidence that FLT3L produced by GREM1 FRCs maintains lymph node preDCs, cDCs, and plasmacytoid DCs (pDCs). Spatial transcriptomics and cytofluorometry reveal that Grem1 FRC-derived FLT3L supports not only proliferation, but also survival of lymph node cDCs within the interfollicular zone (IFZ). Functionally, loss of Grem1 FRC-derived FLT3L impairs cDC priming of antigen-specific T cell responses. These findings provide key mechanistic insights underlying stromal cell support of DC homeostasis and function.

## Introduction

The lymph node is a highly compartmentalized organ, composed of distinct zones that contain specialized niches to optimize antigen encounter and lymphocyte activation. Fibroblastic reticular cells (FRCs) provide the lymph node with a three-dimensional matrix scaffolding^1,2^ and contribute to immunocyte compartmentalization and migration^3,4^ as well as antigen transport^5^. Depletion of FRCs using CCL19- or FAP-cre mice leads to a loss of this lymph node compartmentalization^6,7^, demonstrating their importance in overall lymph node architecture.

Beyond scaffolding, well established lymph node fibroblast subsets possess spatially restricted functions that also support histological organization, homeostasis and function of immunocytes. For example, follicular dendritic cells (FDCs), that reside in B cell follicles, produce CXCL13 to attract B cells, capture antigens, and help germinal center formation^8,9^. Marginal zone reticular cells (MRCs) reside in the outermost cortex and support subcapsular macrophages via production of RANKL^10,11^. T zone FRCs (TRCs) produce immunocyte-directing chemokine gradients of CCL19 and CCL21^7,12,13^ that position DCs in proximity to naive T cells, thereby increasing potential antigen encounter. TRCs also provide the growth factors, IL-7 and BAFF, for naïve T cell and B cells, respectively, maintaining lymphocyte homeostasis^6,14^.

Expanding on the framework of lymph node fibroblast subsets, the use of single-cell RNA-Sequencing (scRNA-seq) allowed identification of additional, previously unseen transcriptional heterogeneity within these broad fibroblast types^15–22^, but functional characterization of the newly identified fibroblast subsets remains unexplored. One particular subset of TRCs described recently is marked by expression of the BMP antagonist *Gremlin 1* (*Grem1*) and is important for the homeostasis of dendritic cells (DCs)^16^. The molecular mechanism underpinning this homeostatic function is unknown.

Conventional dendritic cells (cDCs) are antigen presenting cells that bridge the gap between immune challenge and the adaptive immune response in the lymph node^23^. Lymph node resident cDCs are derived from circulating preDCs that enter lymph nodes via high endothelial venules and subsequently mature into cDCs^24,25^ that reside in the outer cortex prior to challenge^26,27^. DCs require signaling by the growth factor Fms-like tyrosine kinase 3 ligand (FLT3L) for their formation in and maintenance outside of the bone marrow^28–30^, where they are turned over and replaced every 3-6 days^31,32^. cDC homeostasis is maintained in secondary lymphoid organs by both hematopoietic and nonhematopoietic sources of FLT3L^33–35^. However, it is unclear how discrete cellular sources of FLT3L contribute to DC homeostasis within tissues.

Here we show that both human and murine lymph node FRCs express gene programs that contain *Grem1* and *Flt3l* and express a transmembrane domain-containing isoform of FLT3L. Further, using a novel *Flt3l* floxed mouse model, we demonstrate that FLT3L produced by Grem1 FRCs provides a homeostatic growth signal to DCs specifically within the interfollicular zone (IFZ), thereby maintaining optimal resident cDC and pDC numbers in the lymph node. Through advanced computational integration of single-cell and spatial transcriptomics, we identify transcriptional cDC programs controlled by Grem1 FRC-derived FLT3L specifically in the IFZ. Importantly, we demonstrate that maintenance of DCs by Grem1 FRC-derived FLT3L is essential for optimal downstream antigen-specific T cell response following immunization.

## Results

### *Gremlin 1* FRCs express *Flt3l* in human and mouse lymph nodes

In an effort to pinpoint the mechanism by which Grem1 FRCs support lymph node DCs, we applied consensus non-negative matrix factorization (cNMF) to a human lymph node fibroblast scRNA-seq dataset (GSE193449) to identify FRC-specific gene co-expression programs. This analysis led to the identification of 8 major gene programs within human lymph node fibroblasts (Fig. 1a, Extended Data Fig. 1a and Table 1). The FRC program NMF 8 was characterized by expression of *GREM1* (Fig. 1b, c). Notably, *FLT3LG* was also enriched in the *GREM1* program in these FRCs (Fig. 1d). RNA *in situ* hybridization confirmed that *GREM1* and *FLT3LG* transcripts are detected in a spatially colocalized pattern in human lymph nodes (Fig. 1e). Importantly, *GREM1* and *FLT3LG* transcripts were located in close proximity to CD11c/*ITGAX*^+^ DCs (Fig. 1e).

**Fig. 1:**
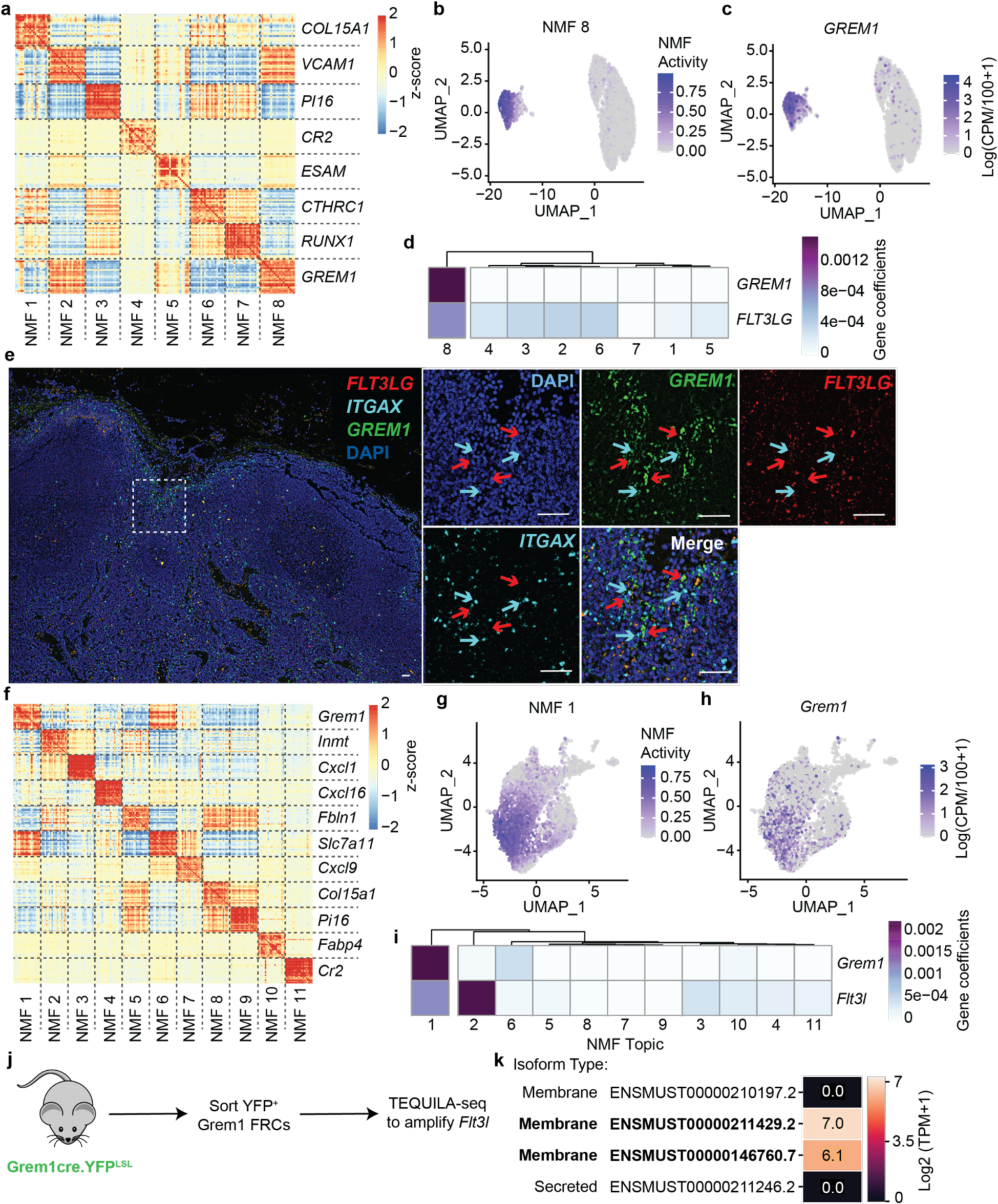
Grem1 FRCs express membrane tethered FLT3LG. **a)** Heatmap of relative (z-score) pairwise Pearson correlation coefficients for top 20 genes for each of the 8 NMF programs derived from cNMF analysis on human lymph node fibroblasts. One representative gene per program is highlighted. **b)** UMAP representation of human lymph node fibroblasts colored by the activity of NMF 8. **c)** UMAP as in (b) colored by *GREM1* expression. Color scaled for normalized expression. **d)** Heatmap of *GREM1* and *FLT3LG* contributions to each NMF program. Columns are grouped by hierarchical clustering. **e)** Representative image (N = 5) of RNAscope from human skin draining lymph nodes show *FLT3LG*^+^ *GREM1*^+^ FRCs (red arrows) near *ITGAX*+ cells (cyan arrows). DAPI (blue), *GREM1* (green), *FLT3LG* (red), *ITGAX* (cyan). Scale bar = 50 um. **f)** Heatmap of relative (z-score) pairwise Pearson correlation coefficients for top 20 genes for each of the 11 NMF programs derived from cNMF analysis on mouse lymph node fibroblasts. **g)** UMAP representation of mouse lymph node fibroblasts colored by the activity of NMF 1. **h)** UMAP as in (g), here colored by *GREM1* expression. **i)** Heatmap of *GREM1* and *FLT3LG* contributions to each NMF program. Columns are grouped using hierarchical clustering. **j)** Schematic of TEQUILA-seq experiment. YFP+ FRCs from Grem1cre.YFP^LSL^ mice were sorted from pooled lymph nodes prior to TEQUILA-seq of *Flt3l*. **k)** Heatmap visualizing the expression levels of 4 annotated protein coding *Flt3l* isoforms.

Next, we investigated whether a *GREM1*/*FLT3LG-*containing gene program can also be found in FRCs from mouse lymph nodes. cNMF analysis of murine FRC scRNA-seq data (E-MTAB-10197) led to the identification of 11 gene programs (Fig. 1f, Extended Data Fig. 1b and Table 2). Program NMF 1 was enriched in *Grem1* expressing FRCs (Fig. 1 g,h). This analysis also revealed two programs with enrichments of *Flt3l*, including NMF 1 (Fig. 1i). Collectively, these results suggest that FRCs co-expressing *Grem1* and *Flt3l* are present in lymph nodes and reside in proximity to DCs.

*Flt3l* can be expressed as secreted or transmembrane spanning isoforms^36^. To determine the mRNA isoform expressed by Grem1 FRCs, we performed targeted long read sequencing, or TEQUILA-seq^37^, on YFP^+^ Grem1 FRCs sorted from lymph nodes of Grem1.ERT2Cre.LSL.YFP mice^16^ (Fig. 1j). Two of the four known protein coding isoforms of *Flt3l* (ENSMUST00000146760.7 and ENSMUST00000211429.2) were expressed by Grem1 FRCs (Fig. 1k). The isoforms expressed by Grem1 FRCs possess membrane spanning regions, indicating that Grem1 FRCs express membrane associated forms of FLT3L and not the soluble isoforms. These data suggest that *Flt3l* is part of a gene program active in Grem1 FRCs that includes expression of the transmembrane form of *Flt3l*.

### Grem1 FRC-derived Flt3l maintains DC numbers in lymph nodes

Next we sought to determine whether expression of *Flt3l* by Grem1 FRCs is mechanistically required for their function in supporting lymph node DCs^16^. To definitively address this possibility, a FLT3L floxed mouse was created and crossed with our previously described Grem1.ERT2Cre knockin (Grem1^wt/ki^) mouse^16^ to generate a novel mouse model in which *Flt3l* can be inducibly and conditionally knocked out in Grem1 expressing fibroblasts (Fig. 2a).

**Fig. 2:**
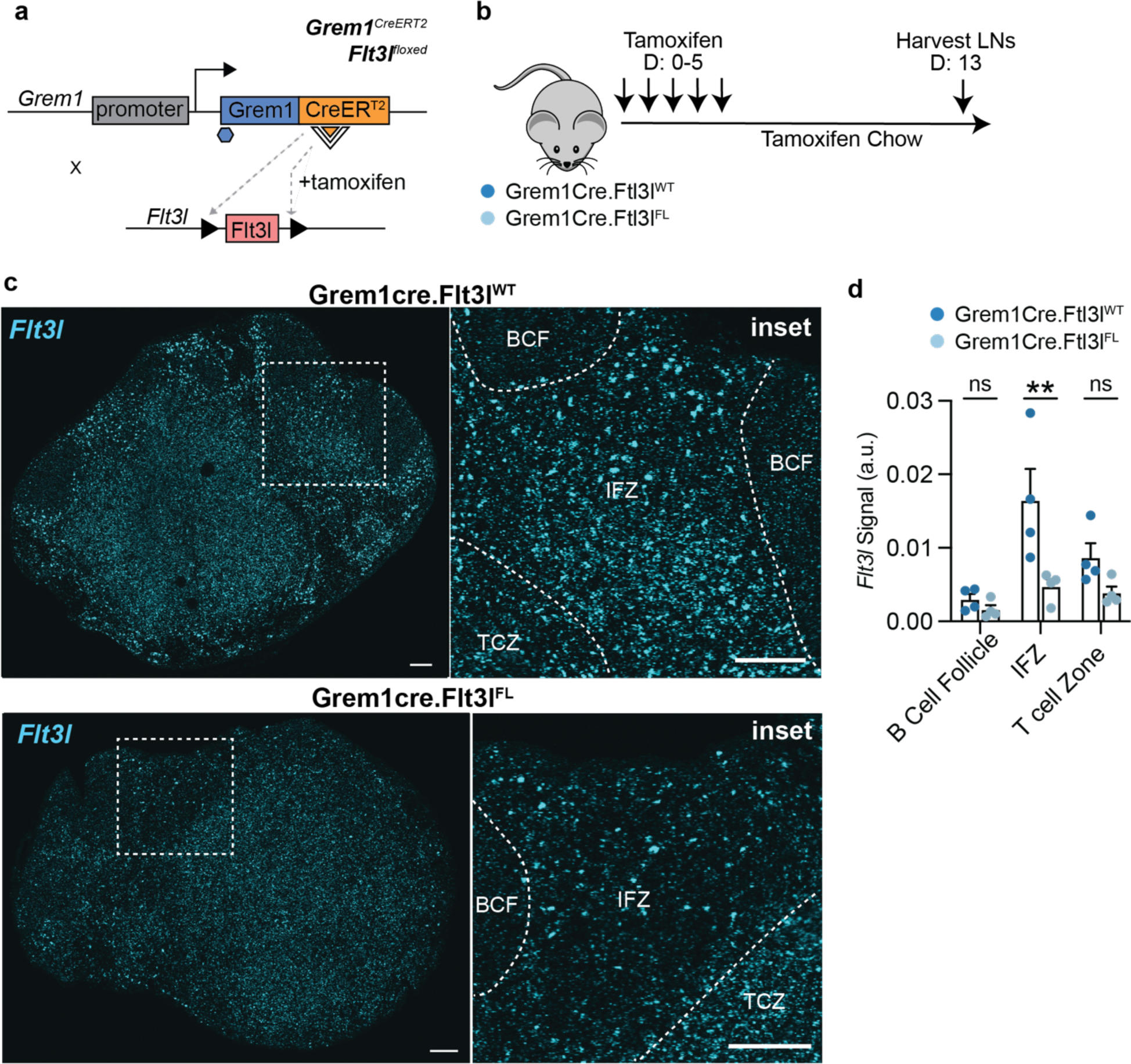
Grem1 FRCs provide an interfollicular zone niche of FLT3L in the lymph node. **a**) Schematic of Grem1cre.Flt3l^FL^ mouse. **b)** Experimental tamoxifen regimen for the Grem1cre.Flt3l^FL^ mouse. **c)** Representative RNAscope images of inguinal lymph nodes from Grem1^wt/ki^Flt3l^fl/fl^ mice and Grem1^wt/ki^Flt3l^wt/wt^ control mice. Inset: Zoomed regions of interest with B cell follicle (BCF), T cell zone (TCZ), and Interfollicular zone (IFZ) annotated. *Flt3l* transcripts (Cyan). Scale bar = 100 um. **d)** Quantification of *Flt3l* signal separated by zone of lymph nodes. Dots represent one image from an individual mouse, bars and error bars represent the mean and standard deviation for each plot. T-test used for statistics. **p<0.01, ns p>0.05.

To ensure sufficient cre activation, Grem1^wt/ki^Flt3l^fl/fl^ (Flt3l-conditional knockout) mice and Grem1^wt/ki^Flt3l^wt/wt^ control mice were placed on a tamoxifen (TAM) regimen in which TAM was administered intraperitoneally for 5 days followed by maintenance on tamoxifen chow for the duration of all functional studies, generally out to days 13-14 (Fig. 2a, b). RNA *in situ* hybridization of inguinal lymph nodes taken from Grem1^wt/ki^Flt3l^wt/wt^ control mice showed a gradient of *Flt3l* expression with highest levels in the interfollicular zone (IFZ), or the area between the T cell zone and B cell follicles, followed by the T cell zone, and lowest levels in the B cell follicle (Fig. 2c, d). Importantly, in Grem1^wt/ki^Flt3l^fl/fl^ mice, *Flt3l* was significantly reduced specifically in the IFZ whereas *Flt3l* was only modestly reduced in the T cell and B cell zones. This pattern of *Flt3l* loss is consistent with the spatial location of Grem1 FRCs and accompanying activity of cre recombinase in the Grem1.ERT2Cre.LSL.YFP genetic mouse model^16^.

Our next step was to evaluate the impact of deleting *Flt3l* in Grem1 FRCs on the cellular composition of lymph nodes by flow cytometry. Total lymph node cells, CD3^+^ T cells and B220^+^ B cells remained unchanged in skin-draining (inguinal) lymph nodes of Grem1^wt/ki^Flt3l^fl/fl^ mice compared to Grem1^wt/ki^Flt3l^wt/wt^ control mice (Fig. 3a-c). In contrast, total DCs (CD11c^+^MHC-II^+^) were significantly reduced in Grem1^wt/ki^Flt3l^fl/fl^ mice (Fig. 3d). Examining DC subsets in Grem1^wt/ki^Flt3l^fl/fl^ mice revealed a reduction in resident cDCs (CD11c^+^MHC-II^mid^, Fig. 3e, f) and in plasmacytoid DCs (pDCs) (CD317^+^, Extended Data Fig. 2a) compared with Grem1^wt/ki^Flt3l^wt/wt^ controls, whereas the number of migratory cDCs (MHC-II^hi^CD11c^+^) remained unchanged (Fig. 3e, g). Similar changes in resident cDCs were observed in cervical and mesenteric lymph nodes (Extended Data Fig. 2b-e) of Grem1^wt/ki^Flt3l^fl/fl^ mice compared with Grem1^wt/ki^Flt3l^wt/wt^ control mice, whereas no changes in DC numbers were observed in skin and spleen (Extended Data Figs. 2f-k). Taken together, these results suggest that Grem1 FRCs provide a source of FLT3L that significantly and preferentially impacts resident cDCs and pDCs within lymph nodes.

**Fig. 3:**
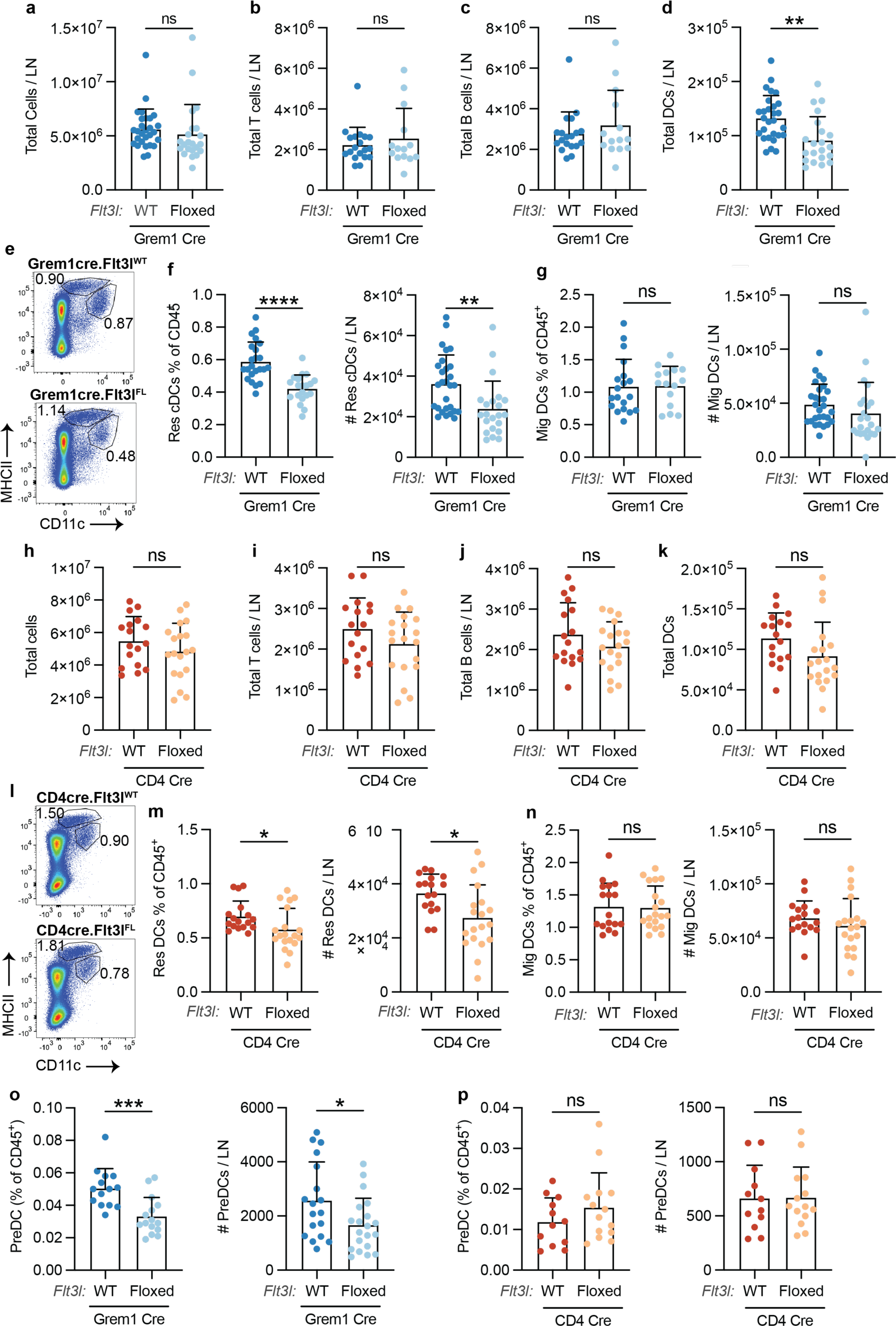
Grem1 FRC niche of FLT3L maintains DC homeostasis in lymph nodes. **a-d)** Quantification of flow cytometry of total cells (a), T cells, (b), B cells (c), and Total DCs (d) in inguinal lymph nodes from Grem1^wt/ki^Flt3l^fl/fl^ mice or Grem1^wt/ki^Flt3l^wt/wt^ control mice. **e)** Representative flow cytometry plots of migratory DC and resident DC populations in inguinal lymph nodes of Grem1^wt/ki^Flt3l^fl/fl^ mice and Grem1^wt/ki^Flt3l^wt/wt^ control mice. Gated on live CD45^+^ single cells. **f,g)** Quantification of flow cytometry (as percent of CD45^+^ cells and total number) of resident cDCs (f) and migratory DCs (g) Grem1cre.Flt3l^FL^ mice or controls. **h-k)** Quantification of flow cytometry of total cells (h), T cells (i), B cells (j), and Total DCs (k) in inguinal lymph nodes from CD4^pos^Flt3l^fl/fl^ mice or controls. **l)** Representative flow cytometry plots of migratory DC and resident DC populations in inguinal lymph nodes of CD4^pos^Flt3l^fl/fl^ mice or controls. Gated on live CD45^+^ single cells. **m,n)** Quantification of flow cytometry (as percent of CD45^+^ cells and total number) of resident cDCs (m) and migratory DCs (n) in CD4^pos^Flt3l^fl/fl^ mice or controls. **o)** Quantification of flow cytometry of live PreDCs (CD45^+^CD11c^+^CD3e^-^CD19^-^I/A-I/B^-^Sirpa^-^) in inguinal lymph nodes from Grem1cre.Flt3l^FL^ mice or controls as percent of CD45^+^ and total number. **p)** Quantification of flow cytometry of preDCs in inguinal lymph nodes from CD4cre.Flt3l^FL^ mice or controls as percent of CD45^+^ and total number. Dots represent individual mice from at least 3 individual experiments, bars and error bars represent the mean and standard deviation for each plot. T-test used for statistics. * p<0.05, **p<0.01, ****p<0.0001, ns p>0.05.

### Limited contribution of T cell-derived FLT3L to homeostasis of lymph node DCs

Previous adoptive transfer studies have shown that T cells represent a key source of FLT3L within secondary lymphoid organs during infection^35^. To what extent T cells contribute to DC homeostasis in lymphoid organs has been unclear. We observed that CD4^+^ and CD8^+^ T cells express transmembrane isoforms of *Flt3l* in non-inflamed lymph nodes (Extended Data Fig. 3a), suggesting that they may also support DC homeostasis in non-reactive lymph nodes.

To determine whether T cell-derived FLT3L affects homeostasis of lymph node DCs, T cell conditional *Flt3l* knockout mice were generated by crossing the *Flt3l*.floxed mouse with a constitutively active CD4 cre mouse (CD4^pos^Flt3l^fl/fl^ mice, Extended Data Fig. 3b). Total cells, CD3+ T cells, B220+ B cells, and surprisingly, total DCs in lymph nodes (Fig. 3h-k) were unaffected in CD4^pos^Flt3l^fl/fl^ mice compared with CD4^pos^Flt3l^wt/wt^ controls. Nonetheless, an assessment of DC subsets in lymph nodes showed that resident cDC numbers were reduced in CD4^pos^Flt3l^fl/fl^ mice (Fig. 3l, m) whereas migratory cDC and pDC numbers were unaffected (Fig. 3l, n; Extended Data Fig. 3c). In spleens of CD4^pos^Flt3l^fl/fl^ mice, total cells, CD3^+^ T cells, B220^+^ B cells, and cDCs were normal compared with CD4^pos^Flt3l^wt/wt^ control mice (Extended Data Fig. 3d-h). These results suggest that T cell derived FLT3L supports resident cDCs in lymph nodes, but is dispensable for the homeostasis of other lymph node DCs as well as all splenic DCs.

### Grem1 FRC-derived FLT3L maintains preDCs in the lymph node

Given that lymph node resident cDCs arise from circulating preDCs, we assessed whether preDCs were also reduced upon deletion of Grem1 FRC-derived FLT3L. Interestingly, cytofluorimetric analysis showed preDCs (CD45^+^CD11c^+^CD3e^-^CD19^-^MHC-II^-^Sirpa^-^)^38^ were significantly reduced in frequency and number in TAM-treated Grem1^wt/ki^Flt3l^fl/fl^ mice compared with Grem1^wt/ki^Flt3l^wt/wt^ control mice (Fig. 3o). In contrast, preDC numbers were normal in lymph nodes of T cell-specific *Flt3l* conditional knockout mice (Fig. 3p). The decrease in lymph node preDCs of Grem1^wt/ki^Flt3l^fl/fl^ mice could be explained by the absence of their growth factor within the lymph node microenvironment or by a reduction in initial lymph node seeding by circulating preDCs. We reasoned that if preDC presence in the lymph node was reduced due to a broader defect in circulating preDCs, a similar reduction in preDCs should be observed in other tissues. To address this possibility, preDCs were enumerated in spleens of the same Grem1^wt/ki^Flt3l^fl/fl^ mice compared to Grem1^wt/ki^Flt3l^wt/wt^ control mice that were used to evaluate lymph nodes. Importantly, the relative abundance and total numbers of preDCs were equal in spleen of Grem1^wt/ki^Flt3l^fl/fl^ mice compared with Grem1^wt/ki^Flt3l^wt/wt^ controls (Extended Data Fig. 4a, b). This finding suggests that lymphoid organ seeding by circulating preDCs is not altered in mice lacking FLT3L expression in Grem1 FRCs.

### FLT3L derived from Grem1 FRCs promotes proliferation and survival of lymph node cDCs

Our finding that Grem1 FRCs provide a key source of FLT3L to preDCs within the lymph node microenvironment raised the possibility of a homeostatic niche for resident DCs. To profile the transcriptional states of DC subtypes upon removal of this essential growth factor specifically in FRCs, we sorted total DCs (CD45^+^MHC-II^+^CD11c^+^) from lymph node of Grem1^wt/ki^Flt3l^fl/fl^ mice and Grem1^wt/ki^Flt3l^wt/wt^ controls and analyzed them by scRNA-seq. We identified 8 unique clusters of DCs from both Grem1^wt/ki^Flt3l^fl/fl^ mice and Grem1^wt/ki^Flt3l^wt/wt^ control mice composed of two clusters of migratory DCs (c0 and c5, *Ccr7, Ccl5*), conventional type 1 DCs (cDC1, c1, *Xcr1, Clec9a*), transitional DCs (tDCs, c2, *Tcf4, Irf4*), conventional type 2 DCs (cDC2, c3, *Cd209a, Sirpa*), Langerhans cells (LH cells, c4, *H2-M2*, *Cd207a*), proliferating cDCs (c6, *Mki67, Top2a*), and plasmacytoid DCs (pDCs, c7, *Bst2, Siglech*) (Fig. 4a,b; Extended Data Fig. 5a,b and Table 3). Due to their low abundance and low expression of MHC class II, we did not capture preDCs in this dataset.

**Fig. 4:**
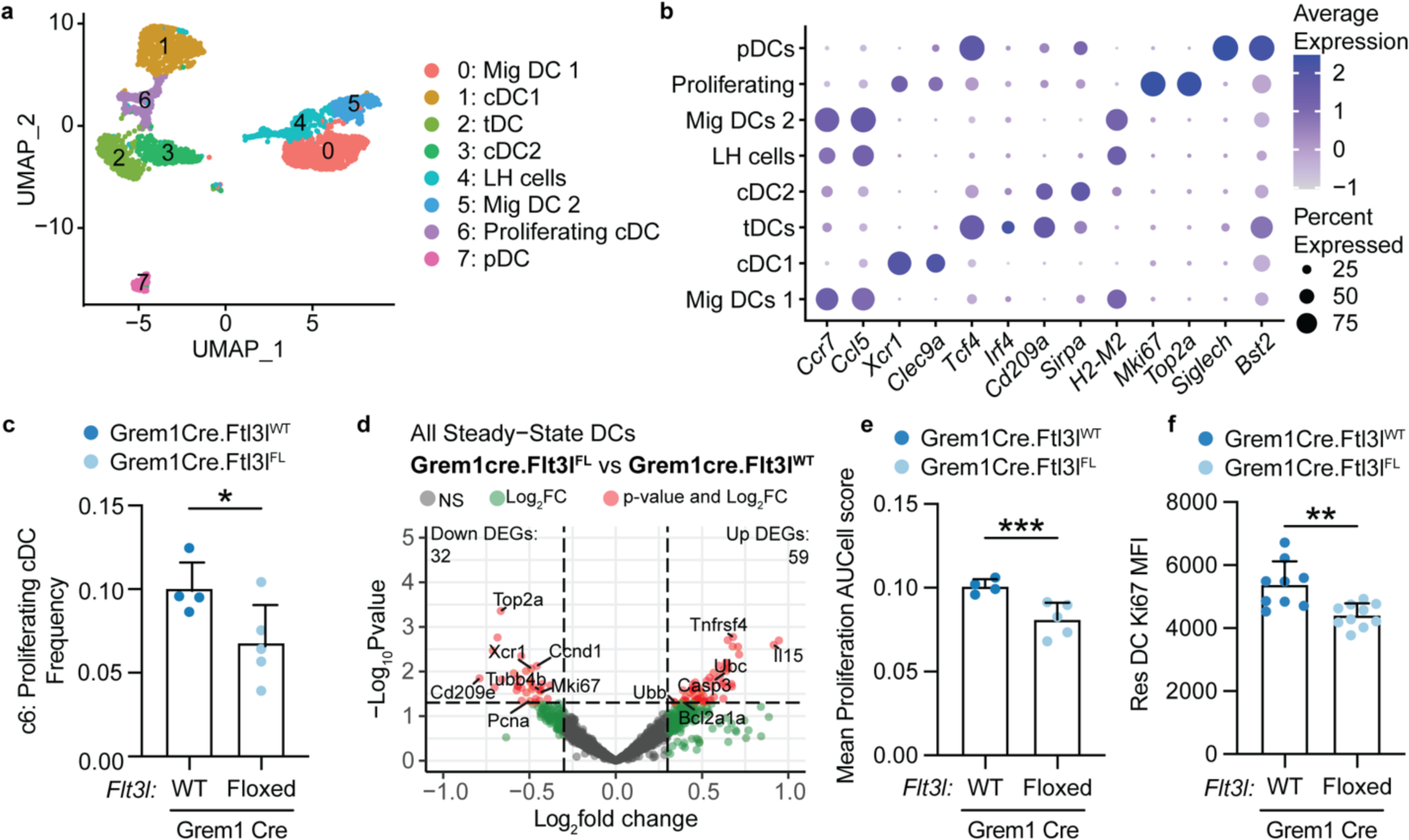
Grem1 FLT3L niche supports homeostatic proliferation of DCs. **a)** UMAP representation of 3,694 cells from sorted DCs (CD45^+^MHC-II^+^CD11c^+^) across Grem1^wt/ki^Flt3l^fl/fl^ mice and Grem1^wt/ki^Flt3l^wt/wt^ control mice, colored by cluster and DC subset annotation. **b)** Dot plot showing the average expression and percent of cells positive for indicated marker genes in each cluster from (a). **c)** Frequency of proliferating cDC cluster (c6) between each condition. Empirical Bayes Moderated T-test was performed using the propeller package. **d)** Volcano plot highlighting log2 fold-change and statistical significance for protein coding genes (dots) between Grem1cre.Flt3l^FL^ vs Grem1cre.Flt3l^WT^ DCs, as identified using pseudobulk analysis in DESeq2. **e)** Mean proliferation gene signature scores comparing Grem1^wt/ki^Flt3l^fl/fl^ mice and Grem1^wt/ki^Flt3l^wt/wt^ control mice. Signature scores computed using the AUCell method. Dots indicate individual mice. **f)** Quantification of a representative experiment for Ki67 MFI in Resident DCs from Grem1^wt/ki^Flt3l^fl/fl^ mice and Grem1^wt/ki^Flt3l^wt/wt^ control mice. Each dot represents an individual mouse, bars indicate mean, error bars indicate standard deviation. T-test was performed unless otherwise specified in the legend. * p<0.05, **p<0.01, ***p<0.001, ns: p>0.05.

Following this transcriptional classification, we investigated how the relative frequency of the 8 DC clusters changed with conditional deletion of *Flt3l* in GREM1 FRCs (Extended Data Fig. 5c). Consistent with cytofluorimetric analysis (Fig.3g), there was no change in frequency of either migratory DC clusters (Extended Data Fig. 5c). However, a significant, 40% reduction in the frequency of proliferating cDCs (c6), was observed in Grem1^wt/ki^Flt3l^fl/fl^ mice compared to Grem1^wt/ki^Flt3l^wt/wt^ controls (Fig. 4c). Consistent with the reduction of proliferating cDCs (c6), pseudobulk differential expression and pathway enrichment analysis revealed a significant reduction in 32 genes associated with cell proliferation, cell division, and E2F target pathways in DCs from Grem1^wt/ki^Flt3l^fl/fl^ compared to Grem1^wt/ki^Flt3l^wt/wt^ controls (Fig. 4d; Extended Data Fig. 5d; Extended Data Table 4). Accordingly, DCs from Grem1^wt/ki^Flt3l^fl/fl^ mice also scored significantly lower for an independent proliferation gene signature (Fig. 4e). We validated these transcriptional findings by flow cytometry, demonstrating decreased Ki67 protein staining in lymph node resident cDCs upon selective depletion of FLT3L in Grem1 FRCs (Fig. 4f). In addition, the 59 up-regulated DEGs in DCs from Grem1^wt/ki^Flt3l^fl/fl^ mice were associated with apoptosis and cell death pathways (Extended Data Fig. 5e), suggesting that Grem1 FRCs also support cDC survival. We did not observe a decrease in frequency of pDCs, however it is possible that this is due to low cellular abundance in our sort. Thus, FLT3L derived from Grem1 FRCs maintains DC numbers within lymph nodes by supporting both proliferation and survival programs.

### Spatially resolved atlas of lymph node cells highlights a DC-supportive niche within the IFZ

We next wanted to ascertain how deletion of Grem1 FRC-derived FLT3L impacts the global transcriptional landscape and architecture of different spatial niches in lymph nodes. To this end, an unbiased, high-resolution single-cell spatial atlas of the mouse lymph node was generated by computationally integrating scRNA-seq (10X Chromium) with spatial transcriptomics (10X Visium) data from paired, replicated samples. For scRNA-seq, we sorted lymphoid (CD45^+^Thy1.2^+^CD19^+^), myeloid (CD45^+^Thy1.2^-^CD19^-^), and stromal (CD45^-^) cells from pooled lymph nodes of 4 Grem1cre.Flt3l^FL^ mice and 4 Grem1cre.Flt3l^WT^ controls, whilst retaining one inguinal lymph node for spatial profiling (Fig. 5a). A total of 24,325 cells were sequenced across six major lineages (Extended Data Fig. 6a-c) that were further annotated into 26 subclusters of cells comprising expected cell populations in the lymph node (Extended Data Fig. 7a-h).

**Fig. 5:**
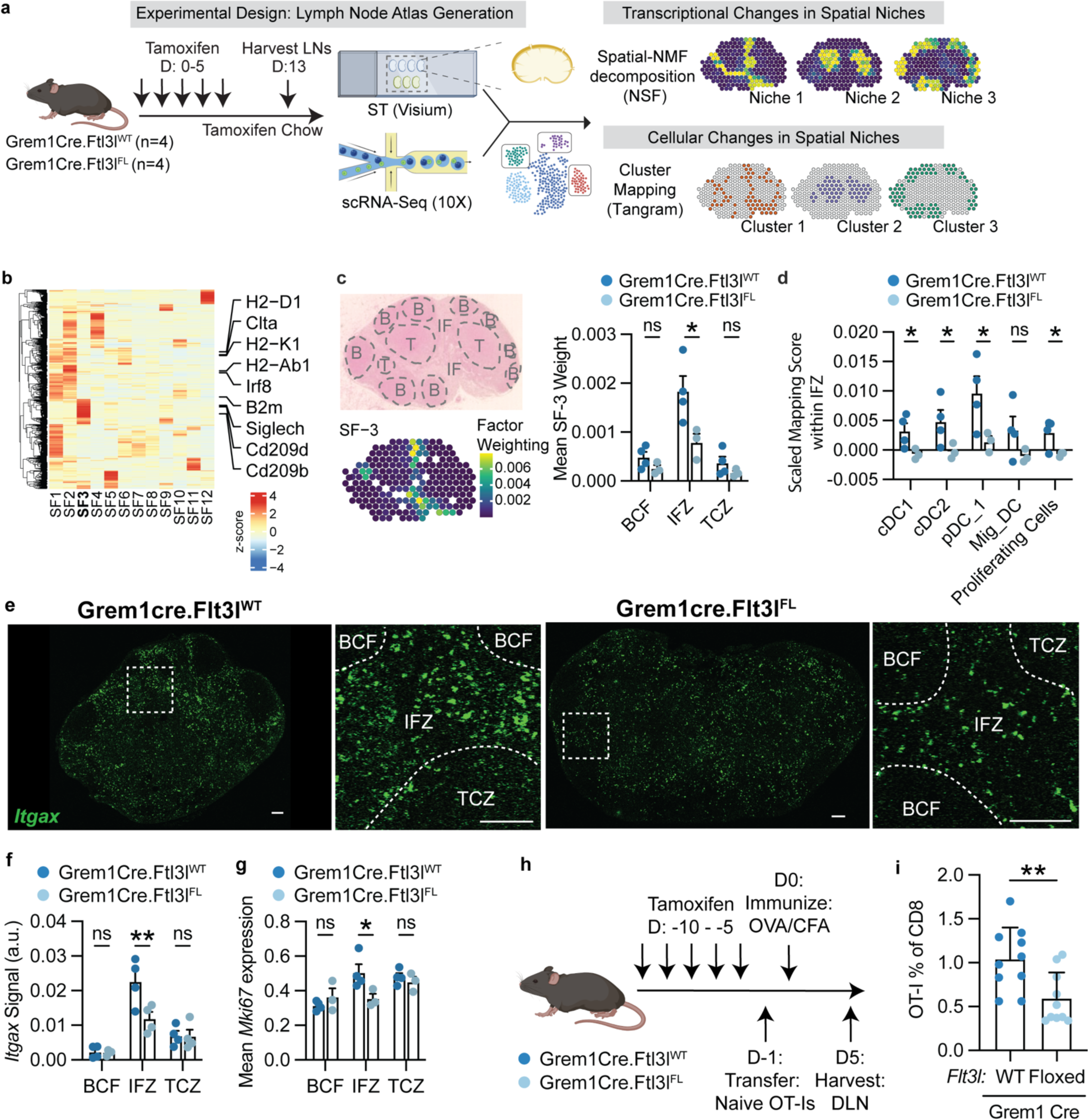
Grem1 FRC FLT3L niche maintains DC homeostasis in IFZ of lymph nodes. **a)** Schematic of the experimental design and analysis strategy of lymph node single-cell RNA-Sequencing and spatial transcriptomics atlas data. **b)** Heatmap of the top-50 genes and their weights for each of the top-12 spatially-aware NMF (NSF) factors. Representative genes enriched in NSF factor SF-3 are highlighted. **c)** Enrichment of spatial niche factor SF-3 in the IFZ of lymph nodes. Representative image of a lymph node (top) from Grem1^wt/ki^Flt3l^wt/wt^ control mouse with manual annotation of spots for B cell zones (B), Interfollicular zone (IF), and T cell zone (T), and a spatial featureplot showing the enrichment of SF-3 in this respective lymph node. Quantification of mean SF-3 weights stratified by lymph node zone. Each dot in the quantification represents the mean factor weight per LN zone of each biological replicate (n=4 WT and n=3 floxed). **d)** Scaled spatial mapping scores, as computed using Tangram, of DC populations, cDC1, cDC2, pDC-1, proliferating cells and migratory DCs, across the IFZ lymph node compartment of the Grem1cre.Flt3l^FL^ mice. Mapping scores (y-axis) are scaled across all lymph nodes. Each dot represents the mean factor weight per LN zone of each biological replicate (n=4 WT and n=3 floxed). **e)** Representative images of RNAscope for *Itgax* (green) from inguinal lymph nodes of a separate cohort of mice. Left: Grem1cre.Flt3l^WT^ mouse, Right: Grem1cre.Flt3l^FL^ mouse. Inset: Zoomed regions of interest with B cell follicle (BCF), T cell zone (TCZ), and Interfollicular zone (IFZ) annotated. **f)** Quantification of *Itgax* RNAscope signal stratified by lymph node zone. **g)** Mean normalized *Mki67* expression from the spatial transcriptomics dataset comparing Grem1cre.Flt3l^WT^ and Grem1cre.Flt3l^FL^ lymph nodes stratified by zone. Each dot represents a biological replicate (n=4 WT and n=3 floxed). **h)** Schematic of the OT-I adoptive transfer and OVA/CFA immunization experiment. **i)** Quantification of OT-I T cells in draining lymph node 5 days post immunization. For all plots, bars represent mean, error bars represent standard deviation. T-test was performed unless otherwise specified in the legend. * p<0.05, ***p<0.001, ns: p>0.05.

For spatial profiling using the Visium assay, all inguinal lymph nodes from Grem1^wt/ki^Flt3l^wt/wt^ and Grem1^wt/ki^Flt3l^fl/fl^ were sectioned and placed into a single reaction square to minimize technical biases in the spatial gene expression comparisons (Fig. 5a, Extended Data Fig. 8a). We manually annotated individual Visium spots across all lymph nodes based on histology for the B cell follicle, IFZ, and T cell zone (Extended Data Fig. 8b). Our annotations were consistent with localization of known marker genes for these compartments (Extended Data Fig. 8c, d).

Since lymph node architecture is organized into well-defined and discrete compartments, we first applied a spatially-aware NMF method^39^ to identify transcriptional programs that align with histological niches. This approach integrates spatial 2-dimensional coordinates as a prior in its matrix factorization model. All 12 spatial NMF factors (SFs) were in strong concordance with the manually annotated lymph node zones (Extended Data Fig. 9a, b and Table 5), demonstrating its ability to generate unbiased spatially-driven transcriptional signatures of different lymph node compartments. To characterize SFs that depend on FLT3L derived from Grem1 FRCs, we compared mean NMF weights between Grem1^wt/ki^Flt3l^fl/fl^ and Grem1^wt/ki^Flt3l^wt/wt^ control lymph nodes for all NMF factors stratified by lymph node zone (Extended Data Fig. 9c). Spatial factor 3 (SF-3) was of particular interest due to its enrichment within the IFZ (Extended Data Fig. 9a,b) and its composition of genes associated with antigen processing and presentation (e.g. *B2m, H2-Ab1, Ciita,* Fig. 5b, Extended Data Fig. 9d). Niche SF-3 was specifically reduced in the IFZ but not in the B cell follicle or T cell zone of Grem1^wt/ki^Flt3l^fl/fl^ lymph nodes relative to controls (Fig. 5c). This finding suggests that FLT3L derived from Grem1 FRCs maintains an antigen presentation program within the IFZ.

Next, we sought to determine the cellular source of the Grem1 FRC FLT3L-dependent antigen presentation gene program within the IFZ and how different DC subsets contribute to it. We mapped all 26 cell clusters across lymphoid, myeloid, and stromal compartments from the whole-lymph node cellular atlas (Extended Data Fig. 6a) onto the Visium lymph node sections using the Tangram method^40^, which aligns paired cellular and spatial transcriptomics datasets. Importantly, mapping a comprehensive single-cell atlas that encompasses a balanced amount of all lymph node cell compartments allowed us to resolve how different cell types are spatially distributed between Grem1^wt/ki^Flt3l^wt/wt^ and Grem1^wt/ki^Flt3l^fl/fl^ lymph nodes (Fig. 5a). Strikingly, cDC1, cDC2, and pDCs clusters showed a decreased mapping score in the IFZ of Grem1^wt/ki^Flt3l^fl/fl^ compared to Grem1^wt/ki^Flt3l^wt/wt^ control lymph nodes (Fig. 5d). Consistent with the cytofluorimetric analysis, which showed no changes in migratory DC presence in the absence of FLT3L expressed by Grem1 FRCs (Fig. 3f), there was no significant change in transcriptional based mapping of migratory DCs to the IFZ (Fig. 5d). We observed similar trends with independent cDC, pDC and migratory DC gene signatures in the IFZ using an orthogonal scoring method which showed reduced cDC and pDC, but not migratory DC scores (Extended Data Fig. 10a). Considering we did not capture preDCs by scRNA-seq due to their very low abundance (Fig. 3o), gene signatures from previously purified preDC clusters^24^ were scored instead (Extended Data Fig. 10b). We observed decreases in preDC signatures that were largely restricted to the IFZ compared to the T cell zone or B cell follicles (Extended Data Fig. 10b). These results are in alignment with our cytofluorimetric data that demonstrated decreased preDCs in lymph nodes of Grem1^wt/ki^Flt3l^fl/fl^ mice (Fig. 3o). Consistently, we found a decrease in the mean expression of the DC marker gene *Itgax* (CD11c) in the IFZ, but not the B cell follicle or T cell zone of Visium sections (Extended Data Fig. 10c). This observation was validated by RNA *in situ* hybridization on a separate cohort of lymph node sections (Fig. 5e, f). Our findings demonstrate that FLT3L specifically derived from Grem1 FRCs maintains preDCs, cDCs, and pDCs (Fig. 3) within the IFZ of lymph nodes, and points to DCs as the cellular source of the IFZ-specific antigen presentation gene program under control of Grem1 FRC-derived FLT3L.

Single-cell RNA sequencing analysis from sorted DCs identified a proliferation program that is supported by Grem1 FRC-derived FLT3L (Fig. 4c-h). Thus, we hypothesized that this program might be actively maintained by DC encounter with FLT3L within the IFZ. Mapping of whole-lymph node scRNA-seq atlas to Visium sections identified a significant reduction in the mapping score of the proliferating cell cluster in the IFZ of Grem1^wt/ki^Flt3l^fl/fl^ mice compared with Grem1^wt/ki^Flt3l^wt/wt^ controls (Fig. 5d). Furthermore, mean expression of the proliferation marker gene *Mki67* was significantly reduced only within the IFZ (Fig. 5g).

Taken together, we performed unbiased spatially-aware gene module identification via NSF and applied Tangram for the mapping of single-cell reference data to a novel LN spatial transcriptomics atlas. Our integrative analysis of results from multiple un-/supervised approaches allowed us to identify previously uncharacterized, lymph node niche-specific transcriptomic changes induced by genetic deletion of FLT3L expression in Grem1 FRCs. The analysis supports a role for Grem1 FRC-derived FLT3L in providing a homeostatic niche for resident DCs within the IFZ.

### Grem1 FRC-derived FLT3L impacts the ability of cDCs to prime antigen-specific T cell responses

To test whether Grem1 FRC-derived FLT3L affects the ability of cDCs to prime antigen-specific T cell responses, we took an adoptive T cell transfer approach. Due to the ability of resident cDCs to present antigens in the draining lymph node following injection^26^, we utilized a subcutaneous immunization model with ovalbumin (OVA). Following TAM administration, naive congenically marked Thy1.1^+^ OT-I T cells with OVA-specific T cell receptors were adoptively transferred into Grem1^wt/ki^Flt3l^fl/fl^ mice and Grem1^wt/ki^Flt3l^wt/wt^ control mice one day prior to immunization with OVA and Complete Freund’s Adjuvant (CFA, Fig. 5h). Using cytofluorimetric analysis, we observed a marked reduction in the frequency of Thy1.1^+^ OT-I T cells 5 days post vaccination in vaccine-draining lymph nodes of Grem1^wt/ki^Flt3l^fl/fl^ mice compared to Grem1^wt/ki^Flt3l^wt/wt^ controls (Fig. 5i). Altogether these data suggest that alterations to lymph node cDC homeostasis can negatively impact priming of antigen-specific T cell responses.

## Discussion

FRCs are transcriptionally and spatially heterogeneous, providing compartmentalization and support for lymph node immunocytes. However, despite significant advances in understanding the transcriptional diversity of lymph node fibroblasts, the field still lacks a comprehensive understanding of the functional roles attributed to this diversity. Here, we profiled a novel genetic mouse model with unbiased single-cell and spatial technologies to define the mechanism by which Grem1 FRCs support DCs. Our data suggest that Grem1 FRCs, which represent a spatially restricted, minor cell population of roughly 1,000 to 2,000 cells per lymph node^16^, provide an essential source of FLT3L for DCs in lymph nodes. In contrast, T cells, which comprise approximately 1000-fold (2,000,000) more cells per lymph node (Fig. 3b), only play a modest role in supporting DC homeostasis in lymph nodes via provision of FLT3L, even when they lack this growth factor throughout development and adulthood. FRCs with high levels of NMF 2 and *Inmt*, may also pose another source of this key growth factor due to their *Flt3l* enrichment. Notably, these FRCs may influence DC homeostasis in the medulla^17^. However, given the role of Grem1 FRCs in DC homeostasis^16^ and the available tools for in vivo assessment, we were unable to explore this source of *Flt3l* more definitively, but its role in DC homeostasis should be explored further.

We propose that the spatial distribution of Grem1 FRCs contributes to their critical role in maintaining DC numbers in lymph nodes. Our findings highlight that Grem1 FRCs, a geographically confined cell population, provide an essential source of FLT3L for preDCs, cDCs, and pDCs in the lymph node IFZ. Consistent with this notion, preDCs^24^, cDCs^26,27^, and pDCs^41^ reside in areas in the lymph node that are similarly populated by Grem1 FRCs^16^, but not T cells. This restricted localization in the IFZ might explain why Grem1 FRC but not T cell sources of FLT3L impact preDCs and pDC numbers. Our data suggest that during their developmental trajectory, DCs require FLT3L signaling from different cellular sources based on their niche. Initially, Grem1 FRCs support preDCs and immature cDCs in the interfollicular zone, while mature cDCs receive FLT3L support from T cells in the T cell zone. Thus, we propose a new function of Grem1 FRCs in which they nurture cDCs in the IFZ at steady-state, readying them for mobilization during immune challenge. Interestingly, neither the Grem1 FRC nor T cell sources of FLT3L impacted migratory DCs, suggesting that migratory DCs are either not in contact with these sources or do not require a source of FLT3L in the lymph node for their homeostasis.

Our findings reveal that Grem1 FRCs express a transmembrane domain containing RNA isoform of *Flt3l*, indicating the potential for direct cellular interaction, requiring cohabitation in the same areas. Interestingly, membrane-bound FLT3L can be cleaved and shed^42^. It remains unclear whether FLT3L shed from Grem1 FRCs signals outside the interfollicular zone or if other sources of soluble FLT3L contribute to DC homeostasis in the lymph node. Exploring the differences in signaling and function between these FLT3L forms could enhance FLT3L-based therapies.

Our approach to understanding the mechanism by which Grem1 FRCs maintain DC homeostasis in lymph nodes identified that Grem1 FRC-derived FLT3L maintains both proliferation and survival of cDCs. Results from our unbiased combinatorial approach of scRNA-seq and spatial transcriptomics indicate that Grem1 FRC-derived FLT3L maintains a transcriptional proliferation program in cDCs, but not pDCs, within the IFZ of LNs. In addition to promoting proliferation of cDCs, perhaps at the preDC to cDC transition^24^, Grem1 FRC-derived FLT3L also decreases expression of apoptosis related genes suggesting Grem1 FRCs maintain both proliferation and survival of cDCs within the IFZ of the lymph node. Overall, we identify a critical role for IFZ-localized Grem1-expressing FRCs in providing FLT3L to lymph node DCs, supporting their proliferation and survival. This work underscores the importance of spatially organized and transcriptionally distinct fibroblastic subsets in creating essential niches for immunocyte development and function.

## Methods

### Mouse models

The Flt3l conditional knockout (Flt3l fl) and Grem1 CreERT2 alleles were generated at Genentech using C57BL/6N ES (C2) cells and established methods. To obtain *Flt3l* fl, exons 2-5 were flanked by loxP sites (floxed), the floxed region corresponding to GRCm38/mm10 chr7:45,133,687-45,135,895. A FRT-flanked Neo cassette used for positive selection in the ES cells was excised using adeno-FLP prior to microinjection, leaving behind a single FRT site. The Grem1 CreERT2, and CD4 Cre mice were maintained in house at Genentech. The Grem1 Grem1^wt/ki^Flt3l^fl/fl^ and Grem1^wt/ki^Flt3l^wt/wt^ mice were maintained on a mixed C57BL/6N and C57BL/6J genetic background. Control mice were non-littermate controls. Thy1.1 OT-I mice were bred and housed at Genentech.

Mice were housed at Genentech in specific pathogen free individually ventilated cages using the guidelines of the US National Institutes of Health. Animals were used between 6 and 12 weeks of age. Age-matched animals were used as controls in all experiments. The sample size and experimental replicate numbers are described in Fig. legends. All animal experiments were performed according to protocols approved by the Genentech Institutional Animal Care and Use Committee.

### Antibodies

Antibodies against the following markers were used for FACS analysis: TCRb (H57-597, 1:200), Ly6C (HK1.4, 1:300), Siglec H (551, 1:200), CD317 (927, 1:200), CD19 (6D5, 1:200), Thy1.2 (30-H12, 1:200), B220 (RA3-6B2, 1:200), Ly6G (1A8, 1:200), XCR1 (ZET, 1:200), CD86 (GL1, 1:200), CD45 (30-F11, 1:200), CD11b (M1/70, 1:200), SIRPa (P84, 1:200), I-A/I-E (M5/114.15.2, 1:400), CD11c (HL3, 1:150), FC Block (2.4G2, 1:200), Ki67 (16A8, 1:100).

### Lymph node and spleen digestion, DC isolation, and tissue preparation

For FACS analysis, FRC and DC isolation for scRNA-seq or bulk sequencing, indicated tissues were collected into RPMI supplemented with 2% Fetal Bovine Serum (FBS). Whole lymph nodes or chopped spleens were then transferred into 15 ml conical tubes containing 2 ml digestion buffer (RPMI containing 2% FBS, 0.1mg/ml DNAse I (Roche, #10104159001), 0.2 m/ml collagenase P (Roche, #11249002001), and 0.8 mg/ml Dispase (Life Technologies, #1710504). Tissues were incubated for 15 minutes in 37°C water bath then triterated by repeated pipetting with a wide-O pipet tip approximately 30 times, supernatant was removed, passed through a 70 um filter and collected on ice. 2 ml digestion buffer was added to incompletely digested tissues for another 15 min incubation and repeated for a total of 2 rounds until tissue is completely digested and no particles are visible. The entire solution was passed through a 70 um filter and collected on ice. Spleens were digested similarly with an additional round of digestion and treatment with RBC ACK lysis buffer for 3 min at room temp. Cells were washed with FACS buffer and centrifuged twice. Single cells were resuspended in FACS buffer for downstream experiments.

### FACS

Single cell suspensions were washed with FACS buffer and stained sequentially with Fixable cell viability dye (Invitrogen, L34957) and primary antibodies for 15 min in the dark on ice, with a wash in between.

### RNA *in situ* hybridization

In-situ hybridization (ISH) was performed using the reagents and protocols from Advanced Cell Diagnostics. Briefly, the lymph nodes were fixed in 10% neutral buffered formalin for 24 hours then transferred into 70% ethanol and processed for paraffin embedding. 5 um thick transverse sections were dried at 60°C for 1 hour. Sections were rehydrated with two washes of xylene for 5 minutes, followed by two washes in 100% ethanol, one wash in 95% ethanol, then 90% ethanol for 1 minute each. Next the sections were incubated in hydrogen peroxide, boiled in antigen retrieval buffer and then digested with proteinase for 15 min at 40°C. After digestion, the slides were washed twice with ISH wash buffer and hybridized with probes for 2 hours at 40°C. Following hybridization, amplifications and secondary hybridization steps were completed for each probe (up to three) according to Advanced Cell Diagnostics protocol. After the final amplification, slides were washed and counterstained with DAPI and then mounted for Immunofluorescence imaging.

### Microscopy

High-resolution images of RNA *in situ* hybridization were acquired on a Nikon A1R confocal or Leica SP8 confocal laser scanning microscope. Full tissue section imaging was performed using a Nikon A1R confocal laser scanning microscope using a 20×Plan Apo lens of NA 0.75.

### Quantification of RNA ISH

Whole lymph node images were manually masked into B cell zones, T cell zones, and the interfollicular zone. Within each mask, *Flt3l* and *Itgax* ISH signal was summed and normalized to total mask area.

### RNA Extraction

Total RNA extraction was performed on isolated cells using the RNeasy Micro kit (Qiagen, Cat No. 74004). RNA was quantified using the RNA high sensitivity Qubit assay (Thermo Fisher Scientific, Cat No. Q32852).

### Full-length cDNA Generation

cDNA was generated from ∼20 ng of input RNA using a custom OligodT primer and TSO. Reverse transcription was performed with Maxima H reverse transcriptase (ThermoFisher, Cat No. EP0751). cDNA was amplified using custom primers (listed in Extended Data Table 6). PCR was performed in a 50 µL reaction volume containing 10 µL of the cDNA product, 1 µL each of the primers (10uM), 25 µL of the 2X LongAmp Taq (NEB, Cat No.M0287S) and 13 µL of H20 according to the following program: 94 °C for 3 min, followed by 13 cycles of (94 °C for 30 s, 60 °C for 15s, and 65 °C for 5 minutes) with a final extension at 65 °C for 5 min. Amplified products were SPRI purified with a 0.6X bead ratio and analyzed on a Tapestation using the High Sensitivity D5000 Screentape assay (Agilent, Cat No. 5067-5592) to assess quality.

To remove unwanted template switching artifacts, a mitigation step was performed to enrich full-length molecules. PCR was performed in a 50 µL reaction volume, containing 20 µL of the amplified cDNA product, 1 µL each of the primers (10 µM), 25 µL 2X LongAmp Taq and 3 µL H20 according to the following program: 94 °C for 3 min, followed by 3 cycles of (94 °C for 30 s, 60 °C for 15s, and 65 °C for 5 minutes) with a final extension at 65 °C for 5 min. The full-length cDNA was captured using 15 µL M270 streptavidin beads (Thermo Fisher Scientific, Cat No.65305) that were washed three times with SSPE buffer (150 mM NaCl, 10 mM NaH^2^PO^4^, and 1 mM EDTA) and resuspended in 10 µL of 5X SSPE buffer (750 mM NaCl, 50 mM, NaH^2^PO^4^, and 5 mM EDTA). The amplified cDNA was combined with 10 µL of the washed M270 beads and incubated at room temperature for 15 minutes. After incubation, the cDNA-bead conjugate was washed twice with 1 mL of 1X SSPE. A final wash was performed with 200 µL of 10 mM Tris-HCl (pH 8.0) and the beads bound to the sample were resuspended in 20 µL H20.

Samples were multiplexed into a single pull-down experiment using a custom barcoding strategy. Off-bead amplification was performed using the barcoded primers (Extended Data Table 6) using the same PCR protocol described above. PCR was performed in a 50 µL reaction volume, containing 20 µL of the sample bead conjugate, 1 µL each of the F/R barcoded primers (10 µM) and 3 µL of H20. A total of 4 cycles of PCR was performed. The barcoded samples were pooled and SPRI purified with a 0.6X bead ratio and eluted in 20 µL H20.

### Probe Generation and Pull down

Custom biotinylated probes targeting genes FLT3L, GREM1, and GAPDH were generated from an oligo pool from Integrated DNA Technologies (IDT). Briefly, probes were designed using the xGen Hyb Panel Design tool (IDT) and were 120-bp in length. Probes were designed to cover all annotated exons and UTRs with a 1x tiling density across gene bodies. Oligo pools were amplified and biotinylated using a nickase-induced linear Strand Displacement Amplification (SDA) approach. The SDA reaction was assembled on ice containing 10 ng of the oligo pool from IDT, 5 µL of the 10X NEB Buffer 3.1, 2mM DTT, 0.25 µM of the RC-oligo, 0.4mM dTTP, 0.6mM dATP, 0.6mM dCTP, 0.6mM dGTP and 0.2mM biotin-16-aminoallyl-2’-dUTP (TriLink BIoTechnologies, Cat No. N-5001). The reaction was first incubated at 95°C for 2 minutes and ramped down to 4°C at a rate of 0.1°C/s. Strand extension was performed at 37°C for 10 minutes using 5uM of ssDNA binding protein (T4 Gene 32 Protein, NEB, Cat No. M0300S) and 0.8U/µL of Klenow Fragment (3’-5’ exo-) DNA polymerase (NEB, Cat No. M0212M). Nickase induced linear SDA was performed using 3nM (0.04U/ µL) of the Nt.BSpQI (NEB, Cat No. R0644S) and was incubated for 4 hours at 37°C. The amplified probes were purified with SPRI beads with a 1.8X bead ratio and eluted in TE Buffer. Probes were analyzed on a 2% Agarose gel to confirm size.

Gene targeting was performed following the IDT protocol (Hybridization capture of DNA libraries using xGen Lockdown probes and reagents). In short, 600 ng of the pooled barcoded cDNA was denatured at 95°C for 3 minutes and incubated with 100 ng of the probes at 65°C overnight. Bead washes and gene capture were performed using M-270 streptavidin beads (Invitrogen, Cat No.65306) per the xGen protocol. The sample-bead conjugate was eluted in 20 µL in H20. PCR amplification was performed using custom primers compatible with Oxford Nanopores cDNA PCR sequencing library prep kit (Nanopore Technologies, Cat No. SQK-PCS111) (Extended Data Table 6). It should be noted that this product is no longer available and has been updated as SQK-PCS114. PCR was performed in a 50 µL reaction volume, containing 20 µL of the sample bead conjugate, 1 µL each of the primers (10X_BC_F_PCS111, 10X_BC_R_PCS111), 25 µL 2X LongAmp Taq and 3 µL H20 according to the following program: 94 °C for 3 min, followed by 3 cycles of (94 °C for 30 s, 53 °C for 15s, and 65 °C for 6 minutes) with a final extension at 65 °C for 6 min. The sample was SPRI purified with a 0.7X bead ratio and eluted in 20 µL of H20. The eluted sample was quantified using the Qubit High Sensitivity dsDNA assay (Thermo Fisher Scientific, Cat No. Q32851). PCR was performed in a 50 µL reaction containing 20 ng of the sample, 1 µL of the cDNA primers (cPRM, SQK-PCS111), 25 µL 2X LongAmp Taq and 16 µL H20 according to the following program: 94 °C for 3 min, followed by 3 cycles of (94 °C for 30 s, 62 °C for 15s, and 65 °C for 6 minutes) with a final extension at 65 °C for 10 min. The sample was SPRI purified with a 0.6X bead ratio and eluted in 20 µL EB buffer. The eluted sample was quantified with the Qubit High Sensitivity dsDNA assay and analyzed on a Tapestation prior to final library prep.

### Library Prep and Nanopore Sequencing

25 fmol of the post-captured cDNA was added into the adapter addition step per the SQK-PCS111 protocol. The final libraries were sequenced on a R9.4.1 MinION flow cell (FLO-MIN106D) and ran for 72 hours.

### Long Read Sequencing Analysis

All nanopore sequencing reads were base called with guppy v7.0.5 using the dna_r9.4.1_450bps_hac model. Sample demultiplexing of barcoded reads was performed with a custom script implementing seqkit (v2.1.0) ‘locate’ with a maximum of two mismatches allowed to the barcode whitelist. Demultiplexed reads were ouput into distinct fastq files and aligned independently with minimap2 (v2.17) to the Gencode mouse reference (GRCm39.primary_assembly.genome.fa). Finally, transcript quantification was performed with isoquant.py (v3.3.0) using the Gencode mouse annotation (gencode.vM33.annotation.gtf).

### Single Cell RNA Sequencing

Lymph node tissue was dissociated into single-cell suspensions as described above. Cells from individual samples were simultaneously stained with corresponding mouse TotalSeq-A hashtag index antibodies (Biolegend; B0301-B0308) and a FACS antibody cocktail (PDPN, CD19, Thy1.2, Ly6G, CD45, CD31, MHCII, CD11c). Indexed samples were washed twice with MACS buffer and pooled together into a single tube for further labeling with live-dead enrichment viability dyes (Calcein Violet and 7AAD). Cells were sorted for live stromal cell fractions (Calcein Violet+, 7AAD-, CD45-, CD31+, PDPN+ and CD31-/PDPN-double negative), lymphocyte fractions (7AAD-, CD45+, CD19+ and Thy1.2+) and myeloid fractions (7AAD-, CD45+, CD19- and Thy1.2-). Cells were counted using a haemocytometer and resuspended in MACS buffer at the appropriate cell concentration for single-cell sequencing using the Chromium Single-Cell v2 3.1ʹ Chemistry Library, Gel Bead, Multiplex and Chip Kits (10x Genomics), according to the manufacturer’s protocol. A total of 10,000 cells were targeted per well.

### Sequencing and single-cell data processing

Gene-expression libraries were sequenced on the NovaSeq platform (Illumina) with paired-end sequencing and dual indexing, whilst hashtag libraries were sequenced on the MiSeq platform (Illumina). A total of 26, 8 and 98 cycles were run for Read 1, i7 index and Read 2, respectively. FASTQ files from gene-expression and hashtag libraries were processed using the Cell Ranger Single Cell v.7.1.0 software (10x Genomics). In brief, scRNA-Seq data were processed using CellRanger *count* function (10x Genomics) and mapped to the mouse reference genome mm10. Filtered gene-cell count matrices that were output from the Cell Ranger count pipeline were imported into Seurat v.4.3.0^43^ using R v.3.6.1 and log-normalized using the *NormalizeData* function. Hashtag antibody count matrices were imported into Seurat and individual samples were demultiplexed using the demuxEM v0.1.7^44^ package. Only cells determined as “Singlets” by demuxEM were included for analysis. Dimensional reduction was performed using the *SCTransform* workflow^45^ built within the Seurat package, which was utilized for downstream Principal Component Analysis (*RunPCA*) and clustering based on shared nearest neighbors (*FindNeighbors*) and leiden graph-based clustering using the original Louvain algorithm (*FindClusters*). A total of 30 principal components (default) were used as input for clustering steps and UMAP reduction (*RunUMAP*).

Broad cell types were first annotated using the SingleR package^46^ with the ImmGen reference gene signature database. For finer cell type annotations, 30 PCAs were used and a resolution of 0.7 was used to identify clusters for the myeloid object; resolution = 0.25 used for lymphoid object; resolution = 0.65 used for stromal cell object; and resolution = 0.4 used for the endothelial cell object. For the stromal cells, clusters were annotated for CCL19 TRCs (*Ccl19, Lepr*), Inmt SCs (*Inmt, Ltbp4),* SMC (*Acta2, Myh11*), Pi16+ fibroblasts (*Pi16, Dpp4*), Dpt+ Fibroblasts (*Col1a1, Ly6a)*, Grem1+ TRC *(Grem1, Ccl21a)* and Col15a1+ Fibroblasts (*Col15a1, Cdh11*). For the myeloid cells, clusters were annotated as Monocytes (*Ifitm3, Lyz2*),

Macrophages (c1: *C1qa, Mrc1*; c6: *Zeb2, Agre4*), pDC_1 (c2: *Siglech, Bst2*; c10: *Runx2*), Migratory DC (*Ccr7, Ccl5*), cDC2 (*Cd209a, H2-DMb2*), cDC1 (*Xcr1, Clec9a*), tDC (*Tcf4, Cd209d*), Neutrophils (*S100a8, S100a9*) and Mast/Basophils (c9: *Mcp8, Gata2*; c11: *Mcpt4, Fcer1a*). For lymphoid cells, clusters were annotated as Naive B cells (*Iglc1, Cd79b*), Germinal center B cells (*Cd83, Nr4a1*), Naive CD8 T-cells (*Cd3e, Cd8a*), Pre B cells (*Bank1, Pax5*), Naive CD4 and T-regs (*Ctla4, Icos*), NK cells (*Nkg7, Klrb1c*), Effector CD8 T cells (*Skap1*), Gamma Delta T cells (*Trdc, Cxcr6*), proliferating cells (*Mki67, Top2a*), IFN B cells (*Ifit3, Isg15*), and Plasma cells (*Ighg1, Ighg2c*). For the endothelial and mural cells, clusters were annotated as High endothelial venules (*Flt1, CD34*), Lymphatic endothelial cells (LEC, *Prox1, Ackr4*), Pericytes (*Rgs5, Pdgfrb*), blood endothelial cells (c3: *Gpihbp1, Cd36*; c4, *Edn1, Sema3g*). Calculation and statistical comparisons of cell cluster proportions were performed using the Speckle package^47^. Pseudo-bulk differential expression analysis was performed using DESeq2^48^ and pooling counts across all cells within each replicate. Results were filtered for protein coding genes using abs(log2FC) > 0.3 and basemean expression >100, and p-value < 0.05.

### RNA QC and Tissue Selection

To assess suitability for spatial transcriptomics, RNA quality of several candidate formalin-fixed paraffin-embedded (FFPE) lymph node and spleen tissues was checked. Two 4 µm tissue curls were tested from each candidate tissue block. The RNA was extracted using the FFPE RNeasy kit (Qiagen #73504) and DV200 percentages were obtained using the Agilent TapeStation 4200 (Agilent, Santa Clara, CA) with High Sensitivity RNA reagents (Agilent #5067-5579/5580). Tissues with a DV200% of 30 or higher were selected for the assay. Multi-tissue blocks were constructed in the HP lab by re-embedding the selected spleens (2/block) and lymph nodes (8/block).

### H&E Staining & ROI Selection

Four-micron sections from the newly-constructed FFPE multi-tissue were placed onto charged slides and a standard H&E (hematoxylin & eosin) stain was performed on the first serial section. The H&E slides were imaged via brightfield microscopy with an Olympus VS200 Research Slide Scanner (Olympus Life Science, Waltham, MA). The second serial slide from each block was designated for the CytAssist Visium assay. An H&E stain was performed on the CytAssist slides following the manufacturer’s guidelines (10x Protocol CG000520 Rev B). The H&E slides were imaged via brightfield microscopy with an Olympus VS200 Research Slide Scanner and shared with the study pathologist who identified regions of interest for the Visium assay. Following ROI selection the slides were decoverslipped, destained and decrosslinked according to Protocol CG000520 Rev B.

### CytAssist Spatial Gene Expression Assay & Library Construction

All steps were done according to 10x Protocol CG000495 Rev E. Temperature-regulated incubation steps were performed using a Bio-Rad C1000 Touch Thermal Cycler (Bio-Rad, Hercules, CA).

Destained slides were immediately loaded into CytAssist Cassettes (11 mm² gasket) for processing. Probe hybridization was performed for 16 hours using 10X Visium Mouse WT Probes (10x #2000455/2000456). Following hybridization, probe ligation and elongation steps were performed. A simple eosin stain was performed to enable visualization of the tissue, and the stained slides were loaded two at a time into the CytAssist instrument. CytAssist-enabled probe release deposited the probes onto CytAssist slides with two 11 mm capture areas. The CytAssist run conditions for all slides was 30 minutes at 37°C. After a probe extension step, the probes were eluted from the CytAssist slides and collected into PCR tubes. A pre-amplification reaction was followed by SPRIselect-mediated cleanup (Beckman Coulter #B23319). Sample index PCR was performed using 10x Dual Index Plate TS Set A (10x #1000251) with 13 cycles. A final SPRI cleanup yielded a library for each capture area.

### Library QC

Library concentration was determined using a Qubit 4 Fluorometer (Invitrogen, Waltham, MA). Library quality was assessed using an Agilent TapeStation 4200 with D5000 dsDNA reagents (Agilent #5067-5588/5589). Finally, NGS adapter presence was confirmed using the KAPA Library Quantification Kit for Illumina systems (Roche #07960140001).

### Visium Spatial Transcriptomics data processing

Sequencing Fastq files were demultiplexed and mapped to the mouse reference genome mm10 using the Space Ranger software v.2.1.1 and the Visium mm10 v.1.0-2020-A probe set (10x Genomics). Processed count matrices were processed using the Seurat package (v.4.1.0) for quality control, data normalization, dimensional reduction and clustering. All Visium spots were annotated for different lymph node compartments (B-cell follicle, Interfollicular zone and T-cell zone) and contaminating adipose regions using the Loupe software (10X Genomics). All adipose regions were excluded for downstream analyses.

### Cellular mapping using the Tangram

Cellular mapping was performed using the Tangram method^40^ using annotated single-cell data from animal matched lymph nodes. To understand the distribution of cell types in the lymph node Visium spatial transcriptomics data, we implemented cluster-level mapping using the Tangram method to determine the composition of each cell type within each measurement spot. Inputs for Tangram mapping are the spot-gene matrix (matrix G, its dimension corresponding to the number of measured spots by number of measured genes), and the averaged gene expression matrix for cell types (Matrix C, its dimension being the number of cell types by number of measured genes). A total of 1,650 genes are used for training, derived by combining the top 100 differentially expressed genes across each cell type. The Tangram cluster-level mapping model jointly learns the mapping matrix M (number of cell type by spots) as well as the overall composition of each cell type, Matrix w.

Cluster level mapping scores computed using Tangram were z-score scaled across all WT and Grem1cre.Flt3l^FL^ mice lymph nodes. The mean cluster mapping scores for each lymph node spatial compartment were compared across control and Grem1cre.Flt3l^FL^ groups. The mean mapping score for all spatial spots in each lymph node compartment were used to derive fold-changes (mean of Grem1cre.Flt3l^FL^ lymph nodes minus the mean for control lymph nodes).

### Identify spatial factors using non-negative spatial factorization

Spatial NMF factors were computed on all Visium lymph node spots using a non-negative spatial factorization (NSF) method^39^. Default parameters for the NSF-Hybrid analysis, as recommended by the developers, were used for computing the top 12 spatial factors. A total of 3000 spatially variable features were used for NSF. Mean factor weights per lymph nodes were used to compare factors across control and Grem1cre.Flt3l^FL^ groups. The top 100 genes, ranked by spatial factor weight, were used for gene ontology analysis with the Biological Processes sub-ontology database in the ClusterProfiler package.

### Gene-signature scoring using AUCell

The top-50 genes (ranked by fold change) from preDC^24^, mig_DC, cDC1, cDC2, pDCs and tDCs clusters, as defined using FindAllMarkers function in the Seurat package (v.4.1.0), were used to define gene-signatures. Signatures were then scored across all Visium spots using the AUCell method^49^. AUCell scores were calculated in only the top 5% of genes ranked by log-normalized expression, as recommended by the developers.

### Identification of mouse and human lymph node FRC NMF programs

Mouse FRC scRNA-seq data was processed as described in^16^. We used cNMF (200 iterations) with the raw count matrix of 4,069 FRCs and the 3,000 most variable genes detected in Seurat (*FindVariableFeatures*) as input. From a range for *k* from 5-20, we selected 11 as the preferred solution considering stability and error and removed outliers with euclidean distance >0.2. NMF program scores within cells were normalized to sum to 1.

For the human scRNA-seq dataset (GSE193449), cNMF was run on the raw counts of 4,877 cells including all genes with at least one count in at least 10 cells. The top 4,000 highly variable genes identified in Seurat (*FindVariableFeatures*) were designated as variable genes in cNMF. cNMF was run with 200 iterations and with a number of components from 3 to 21. Eight components were chosen as this had the highest stability. A threshold for euclidean distance of 0.1 was used to remove outliers. NMF program scores within cells were normalized to sum to 1.

## Data & Code availability

Processed data and analysis code is available through the Open Science Framework (OSF) (https://osf.io/ew6vz). No new algorithms were developed for this manuscript. FASTQ sequencing files and processed count matrices for the single-cell RNA sequencing (10X Chromium) and spatial transcriptomics data (10X Visium) will be deposited to the Array Express database upon publication.

## Acknowledgments

We thank our colleagues in Animal Resources for excellent animal husbandry and our colleagues in Molecular Biology for allele generation and genetic analysis. We thank our colleagues in the FACS core at Genentech and the support that they provided for flow sorting. We thank our colleagues in the NGS core that provided support for scRNA-seq. We thank members of the Turley and Muller lab for insightful discussions. This work was supported by Genentech.

---

**Extended Data Fig. 1:**
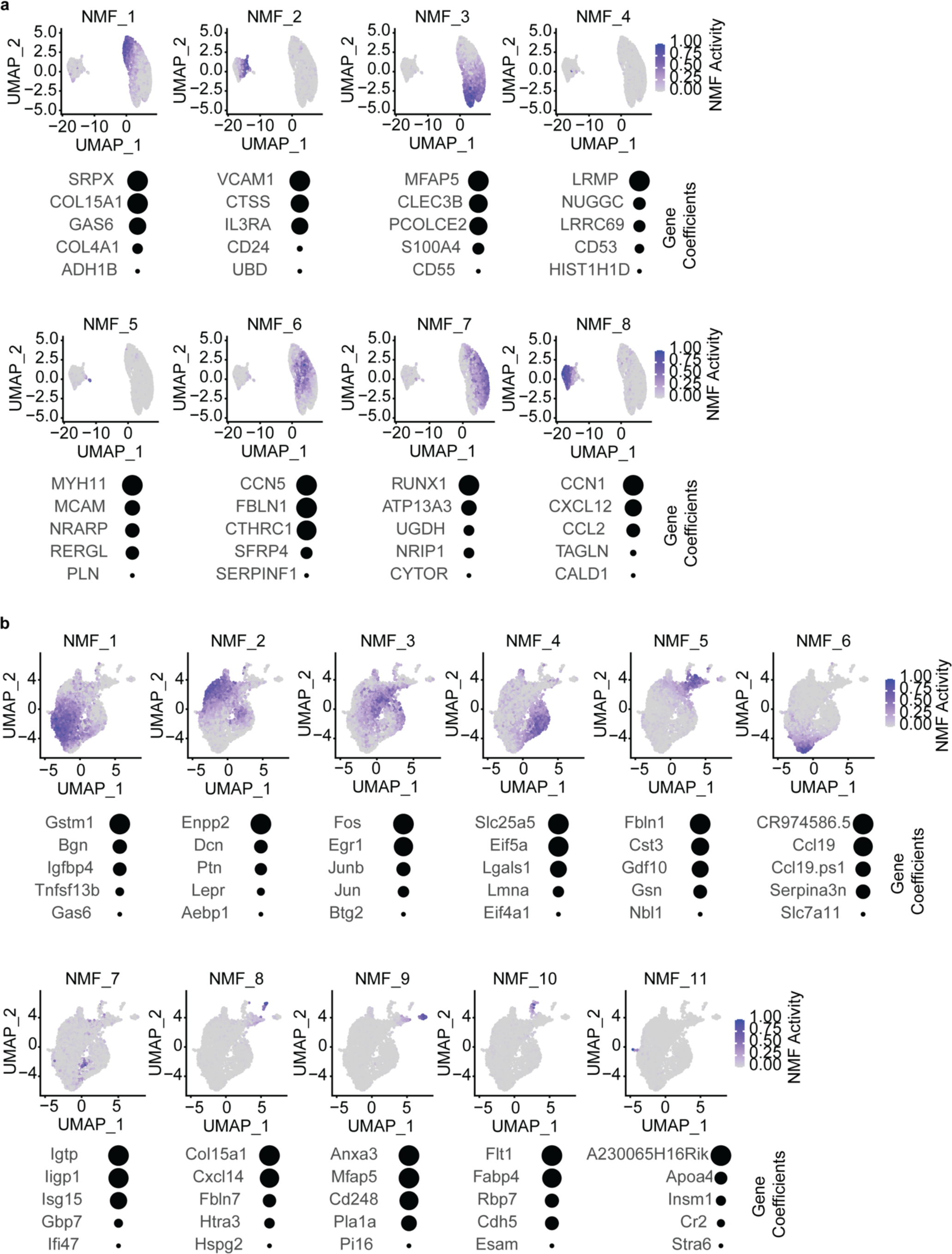
Human and murine lymph node fibroblast NMF programs. **a)** FeaturePlots of each NMF program projected on UMAP of human lymph node fibroblasts (top) and gene coefficients for the top 5 genes for each NMF program (bottom). **b)** FeaturePlots of each NMF program projected on UMAP of murine lymph node fibroblasts (top) and gene coefficients for the top 5 genes for each NMF program (bottom).

**Extended Data Fig. 2:**
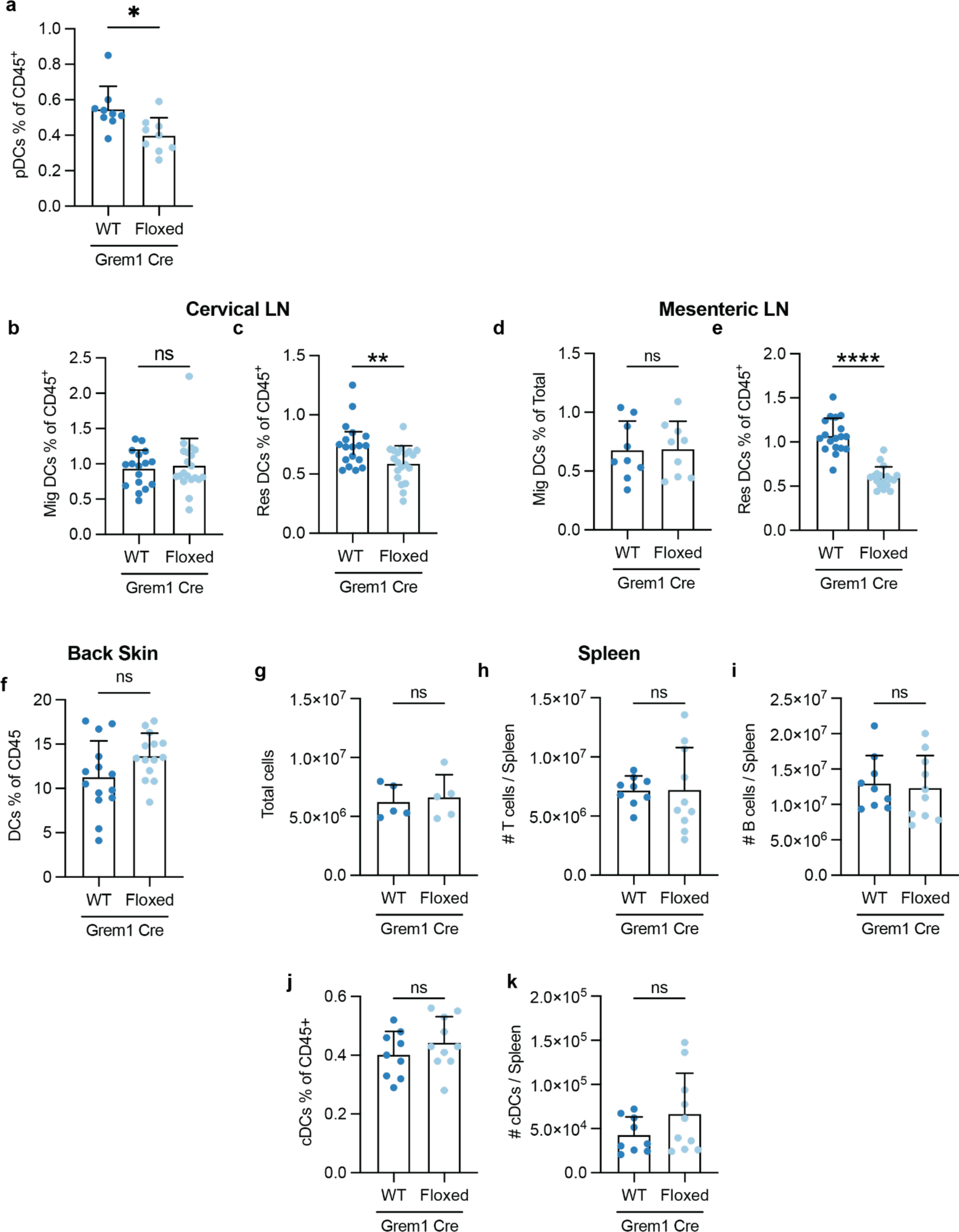
Grem1 FRCs maintain DCs in other lymph nodes, but not spleen or skin. **a)** Quantification of CD317^+^ pDCs in inguinal lymph nodes of Grem1cre.Flt3l^FL^ mice or controls. **b,c)** Quantification of flow cytometry of migratory DCs (b) and Resident DCs (c) from Cervical lymph nodes of Grem1^wt/ki^Flt3l^fl/fl^ mice and Grem1^wt/ki^Flt3l^wt/wt^ control mice. **d,e)** Quantification of flow cytometry of migratory DCs (d) and Resident DCs (e) from Mesenteric lymph nodes of Grem1cre.Flt3l^FL^ mice and controls. **f)** Quantification of flow cytometry of DCs as percent of CD45^+^ cells in the back skin of Grem1cre.Flt3l^FL^ mice and controls. **g-k)** Quantification of flow cytometry of total cells (g), T cells (h), B cells (i) and cDCs (j & k) from spleens of Grem1^wt/ki^Flt3l^fl/fl^ mice and Grem1^wt/ki^Flt3l^wt/wt^ control mice. T-test used for statistics unless otherwise stated in Fig. legend. **p<0.01, ns p>0.05.

**Extended Data Fig. 3:**
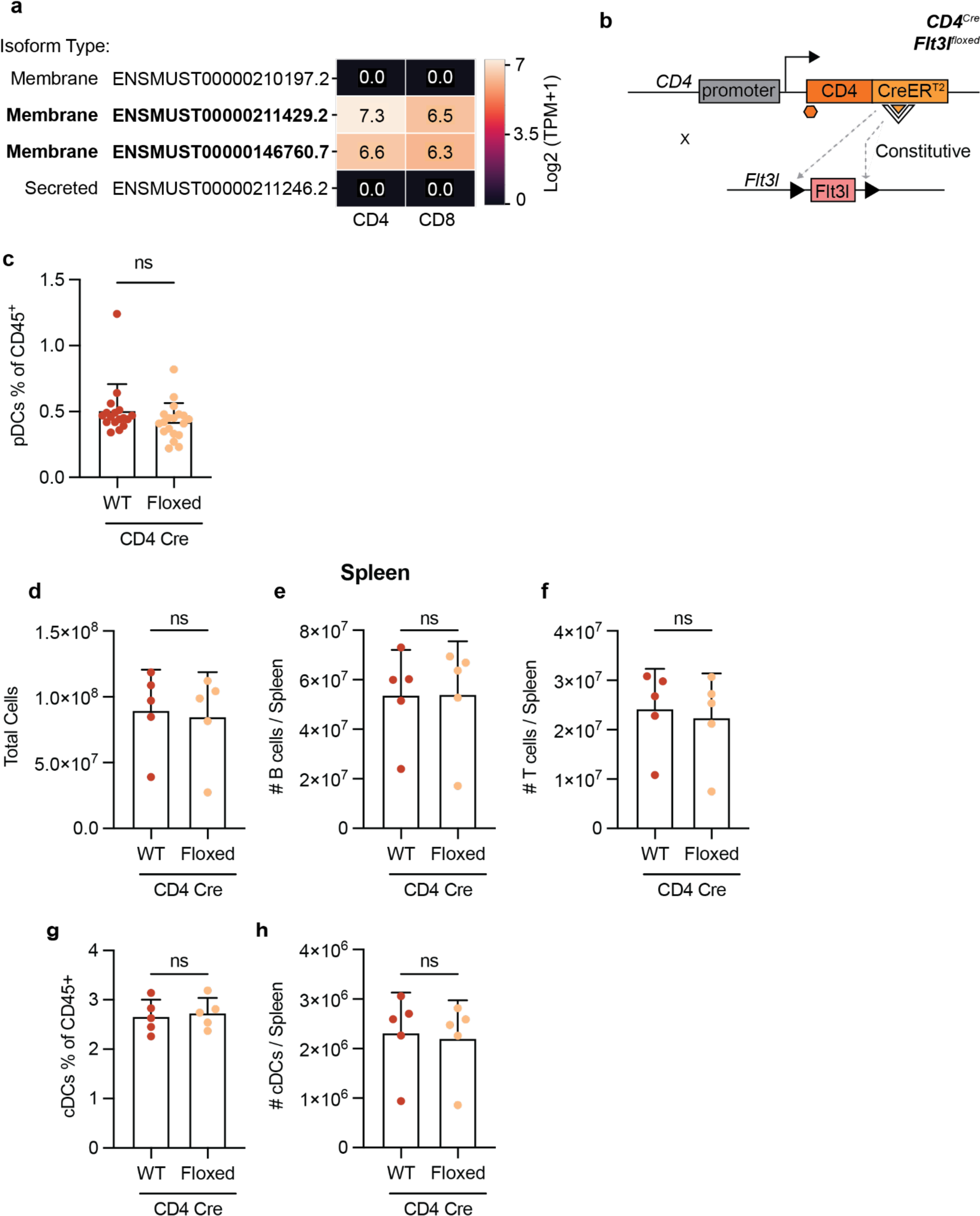
Loss of T cell specific Flt3l has minimal effects on DCs in lymph node and spleen. **a)** TEQUILA-seq from sorted CD4^+^ and CD8^+^ T cells showing the expression of 2 isoforms of *Flt3l* (bolded) that possess membrane spanning regions. **b)** Schematic of CD4^pos^Flt3l^fl/fl^ mice. **c)** Quantification of CD317^+^ pDCs in inguinal lymph nodes of CD4cre.Flt3l^FL^ mice or controls. **d-h)** Quantification of flow cytometry of total cells (d), B cells (e), T cells (f), and cDCs (g & h) from spleens of CD4^pos^Flt3l^fl/fl^ mice. Dots represent individual mice, bars represent mean, error bars represent standard deviation. T-test used for statistics. *p<0.05, ns p>0.05.

**Extended Data Fig. 4:**
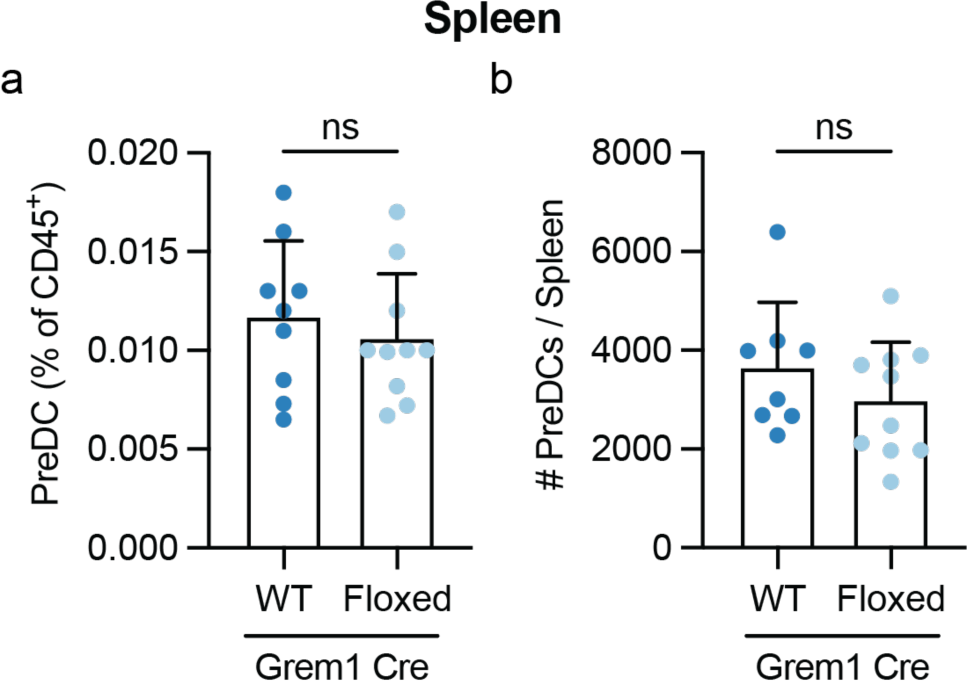
Grem1 FLT3L is dispensable for PreDCs in spleen a,b) Quantification of flow cytometry of PreDCs from spleens of Grem1cre.Flt3l^FL^ mice or controls showing no change in splenic PreDCs as percent of CD45^+^ cells (a) or total number (b) in Grem1^wt/ki^Flt3l^fl/fl^ mice and Grem1^wt/ki^Flt3l^wt/wt^ control mice. Dots represent individual mice from at least 3 individual experiments, bars represent mean, error bars represent standard deviation. T-test used for statistics. ns: p>0.05

**Extended Data Fig. 5:**
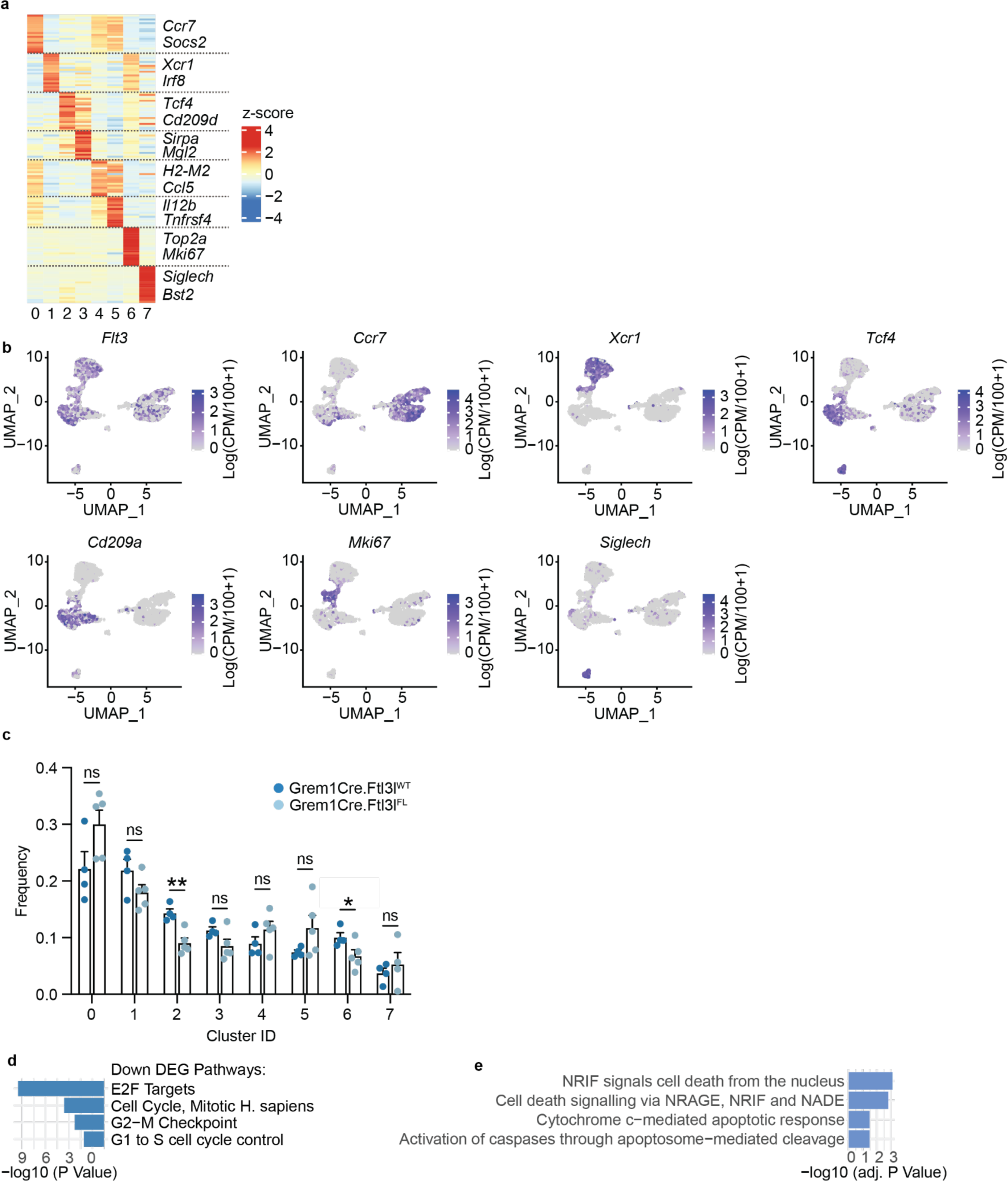
Identification of DC clusters in lymph nodes of Grem1^wt/ki^Flt3l^fl/fl^ mice and Grem1^wt/ki^Flt3l^wt/wt^ control mice. **a)** Heatmap of the top 20 marker genes for each cluster with specific identity genes highlighted from scRNA-seq of DCs (CD45^+^CD11c^+^MHC-II^+^) sorted from Grem1^wt/ki^Flt3l^fl/fl^ mice and Grem1^wt/ki^Flt3l^wt/wt^ control mice from Fig. 4. **b)** FeaturePlots of selected identity marker genes for each cell subset. **c)** Frequencies of each DC cluster between Grem1^wt/ki^Flt3l^fl/fl^ mice and Grem1^wt/ki^Flt3l^wt/wt^ control mice. Dots represent individual hash-tagged mice from the scRNA-seq experiment, bars represent mean, error bars represent standard deviation. Empirical Bayes Moderated T-test was performed using the propeller package. **d)** Gene set enrichment analysis of the 32 down regulated DEG from Fig. 4d. **e)** Gene set enrichment analysis from the 59 up regulated DEGs from Fig. 4d. *p<0.05, **p<0.01, ns: p>0.05.

**Extended Data Fig. 6:**
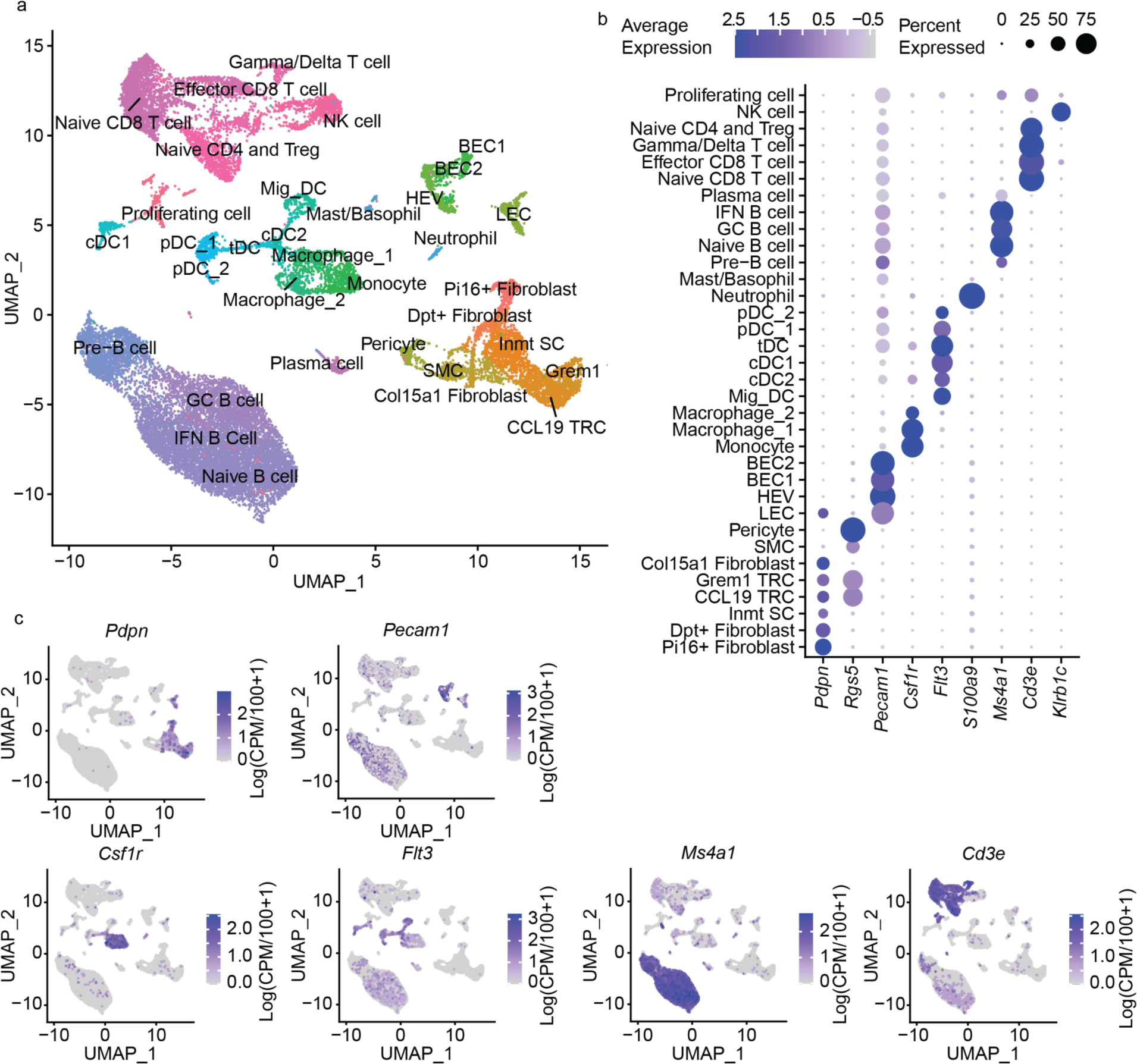
Single cell RNA sequencing atlas of lymph nodes of Grem1^wt/ki^Flt3l^fl/fl^ mice and Grem1^wt/ki^Flt3l^wt/wt^ control mice. **a)** UMAP representation of 24,325 cells sequenced from lymph nodes of Grem1^wt/ki^Flt3l^fl/fl^ mice and Grem1^wt/ki^Flt3l^wt/wt^ control mice. Stromal, myeloid and lymphoid cells are clustered and annotated into 26 specific sub-populations (Described in Extended Data Fig. 7). **b)** Dot plot representing the average expression and abundance of expression of broad cell type marker genes. **c)** FeaturePlots of selected marker genes used to annotate broad cell types from b.

**Extended Data Fig. 7:**
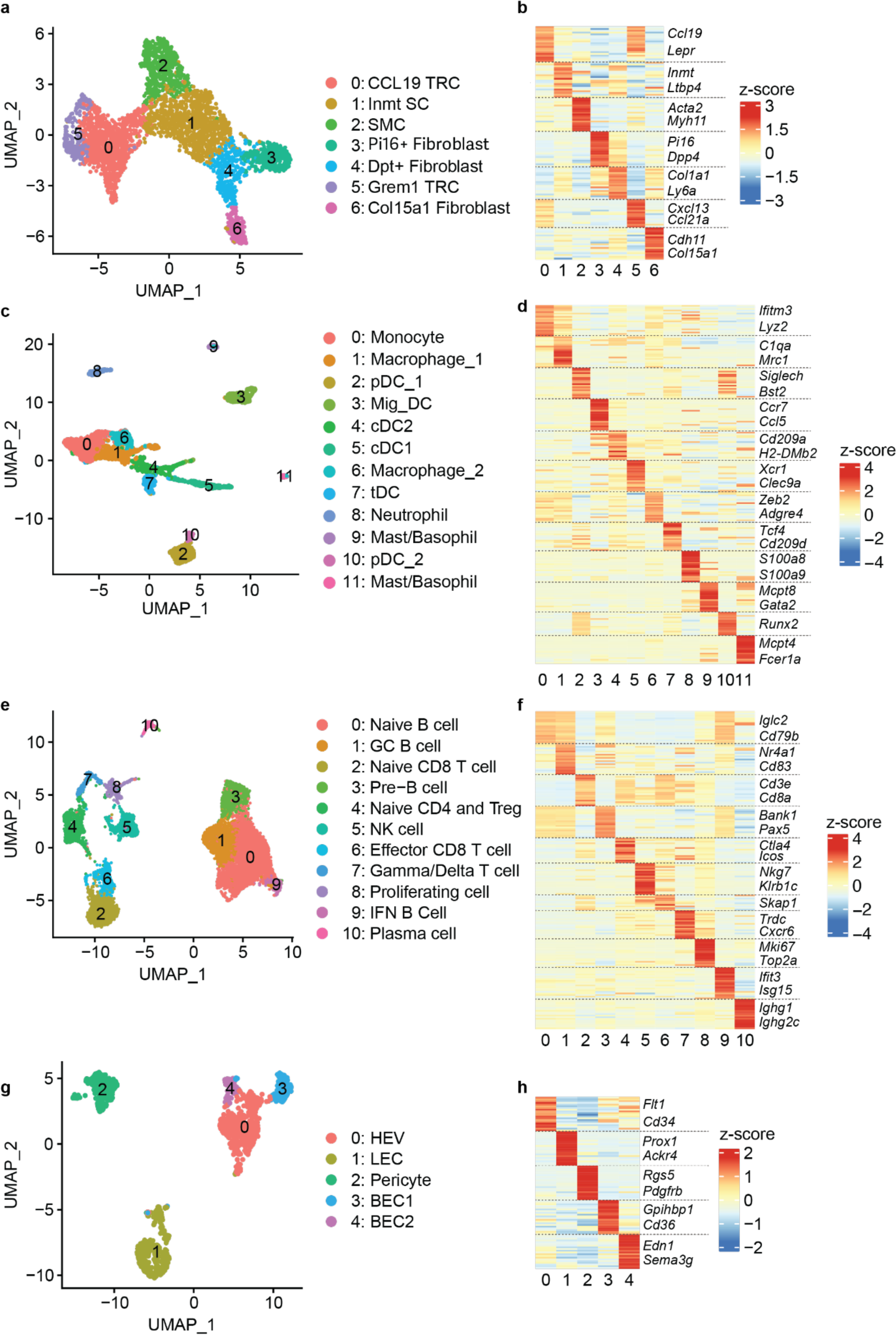
Detailed stromal, myeloid and lymphoid annotations from the lymph node scRNA-seq atlas from Grem1^wt/ki^Flt3l^fl/fl^ mice and Grem1^wt/ki^Flt3l^wt/wt^ control mice. **a)** UMAP clustering of 5,512 cells into distinct clusters of fibroblasts. **b)** Heatmap of top 20 marker genes for each Fibroblast cluster with specific identity genes highlighted. **c)** UMAP clustering of 2,907 cells into distinct clusters of myeloid cells. **d)** Heatmap of top 20 marker genes for each Myeloid cell cluster with specific identity genes highlighted. **e)** UMAP clustering of 16,417 cells into distinct clusters of lymphocytes. **f)** Heatmap of top 20 marker genes for each lymphocyte cluster with specific identity genes highlighted. **g)** UMAP clustering of 1,407 cells into distinct clusters of blood endothelial cells (BECs), lymphatic endothelial cells (LECs) and pericytes. **h)** Heatmap of top 20 marker genes for each cluster of BECs, LECs, and pericytes with specific identity genes highlighted.

**Extended Data Fig. 8:**
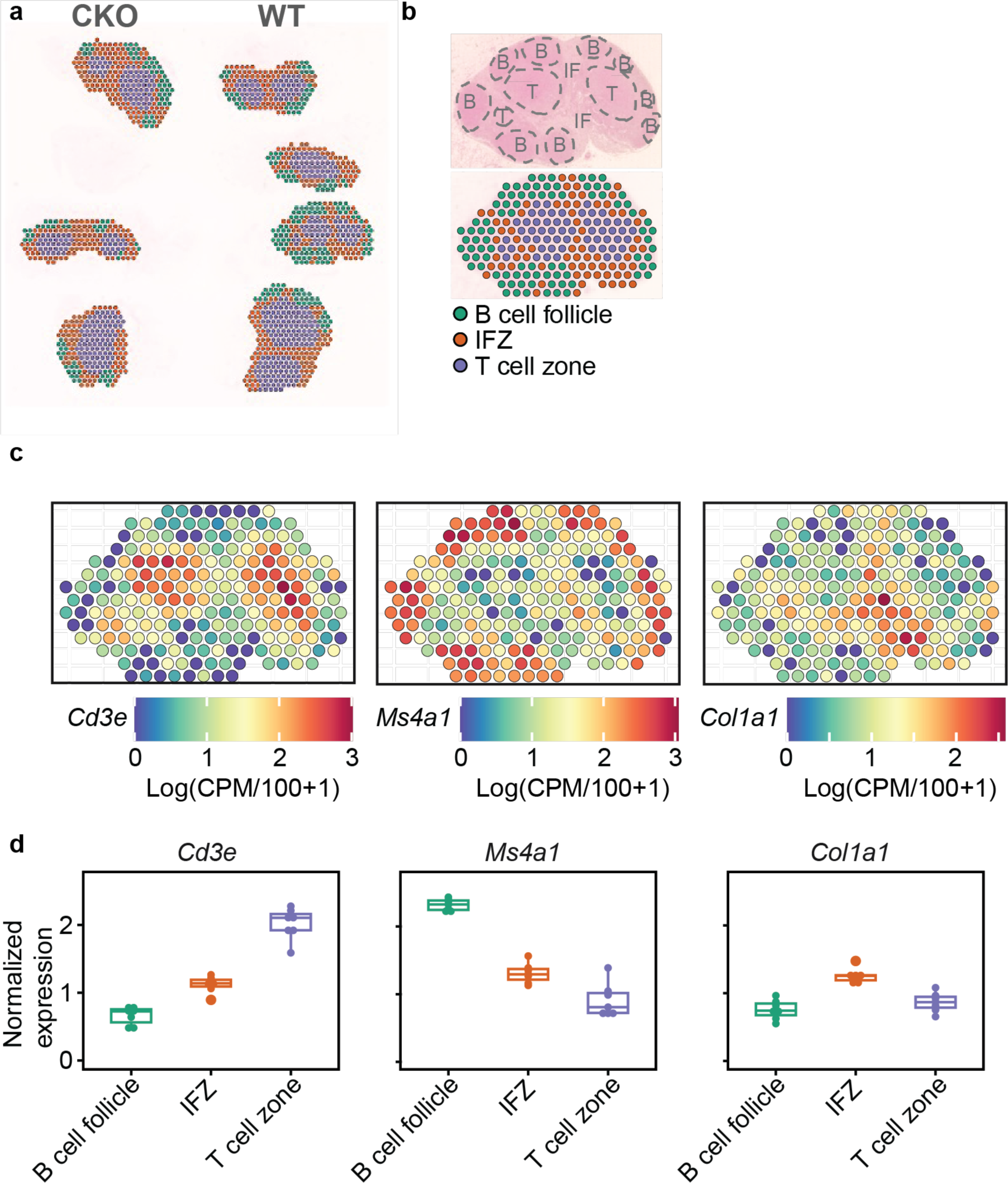
Spatial transcriptomics annotations and gene expression patterns across lymph node zones. **a)** Spatial plot of all lymph samples across Grem1^wt/ki^Flt3l^fl/fl^ mice (n=3) and Grem1^wt/ki^Flt3l^wt/wt^ (n=4) control mice. Spots are annotated by lymph node compartment. **b)** Representative image of a lymph node analyzed using Visium spatial transcriptomics. Manual annotation of spots for B cell zones (B), Interfollicular zone (IF), and T cell zone (T) of lymph nodes are shown. **c)** Expression of *Cd3e* (CD3), *Ms4a1* (CD20) and *Col1a1* (Collagen 1) across the representative lymph node example shown in a) highlighting the specific enrichment across the respective lymph node compartments. **d)** Boxplot of the expression for *Cd3e* (CD3), *Ms4a1* (CD20) and *Col1a1* (Collagen 1) stratified by zone annotation. Each dot represents the mean of individual mouse replicate lymph nodes. Box plots depict the first and third quartiles as the lower and upper bounds, respectively. Whiskers represent 1.5× the interquartile range and the center depicts the median.

**Extended Data Fig. 9:**
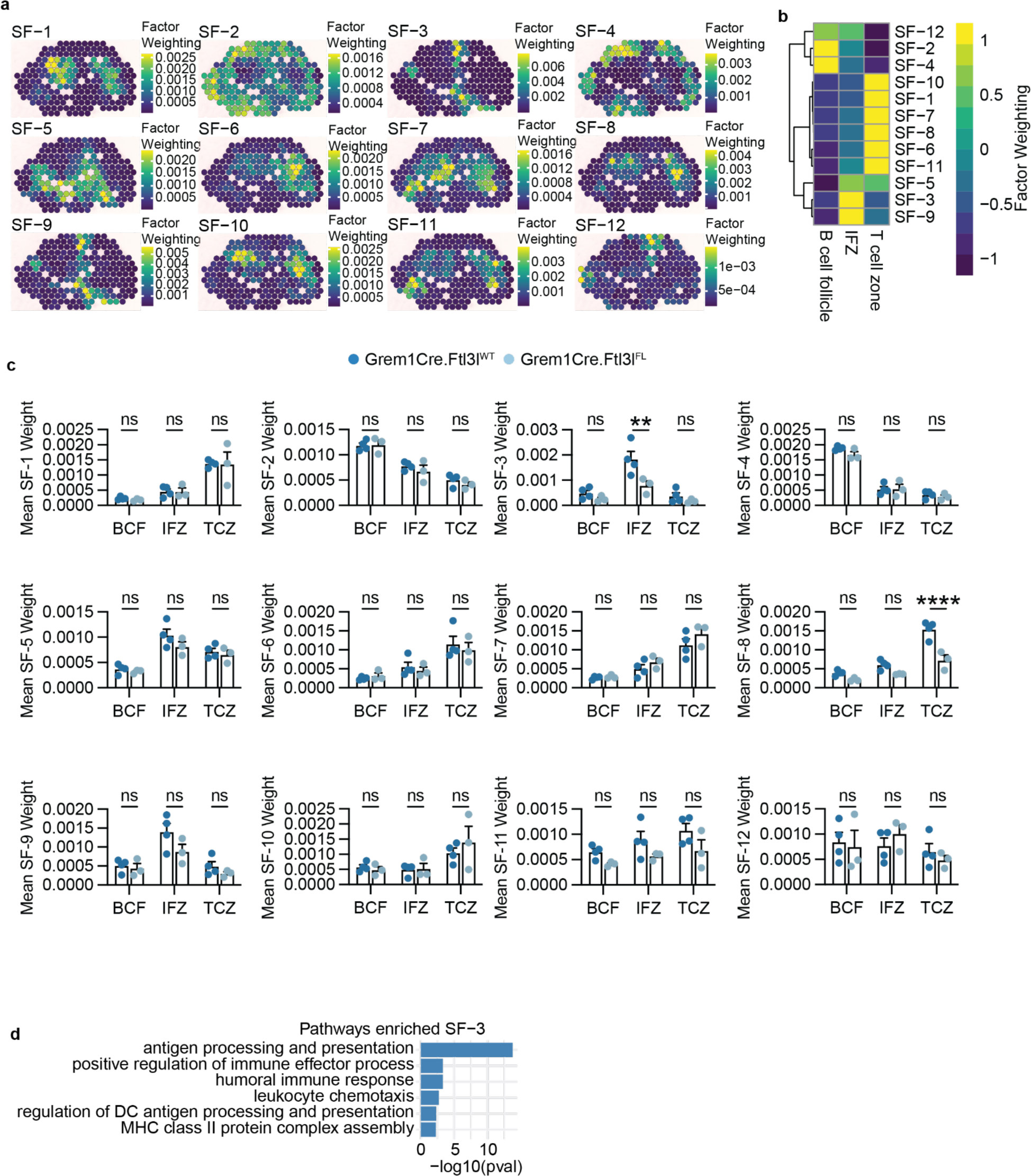
Spatial NMF reveals Grem1 FRC FLT3L maintains spatial programs in distinct areas of lymph nodes from Grem1^wt/ki^Flt3l^fl/fl^ mice and Grem1^wt/ki^Flt3l^wt/wt^ control mice. **a)** Featureplot of the top-12 spatial factors (SFs) in a representative Grem1^wt/ki^Flt3l^wt/wt^ control mouse lymph node section. Factors were computed using spatially-aware non-negative matrix factorization (NSF). **b)** Specific enrichment of NSF factors across different lymph node compartments: B cell follicle, Interfollicular zone (IFZ) and T-cell zone. Heatmap shows the average NSF weight per spot grouped by region. **c)** Mean NSF factor weights (Top-12) compared across specific lymph node compartments between the Grem1^wt/ki^Flt3l^fl/fl^ and Grem1^wt/ki^Flt3l^wt/wt^ control mice. Dots represent individual mouse replicates. Two-Way ANOVA was performed across zones per biological visium replicate. **d)** Gene ontology pathway enrichment of the top 100 genes for NSF factor SF-3. **p<0.01, ****p<0.0001, ns p>0.05.

**Extended Data Fig. 10:**
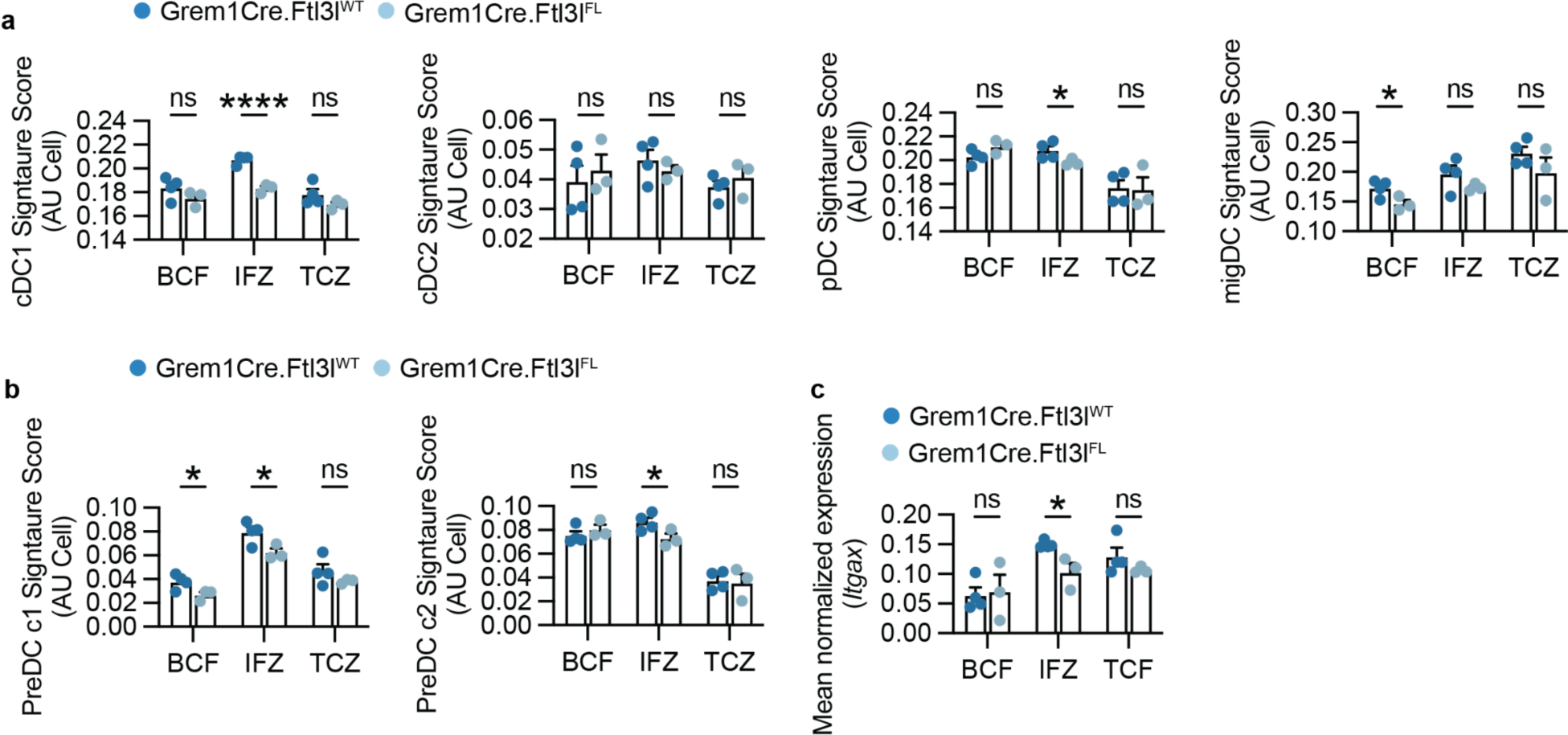
PreDC and DC gene signature scoring across lymph node compartments by spatial transcriptomics. **a)** Signature scores computed using the AUCell method for DC subsets (Fig. 4a), cDC1, cDC2, pDC and migDCs. **b)** Signature scores computed using the AUCell method for PreDC gene signatures from Ugur et al.^24^ **c)** Mean normalized expression of *Itgax* (CD11c) from Visium sections between conditions, stratified by lymph node zone. Dots represent individual mouse Visium sections. T-test was used for statistical analysis. * p<0.05, ****p<0.0001, ns p>0.05.

